# A role for conformational changes in enzyme catalysis

**DOI:** 10.1101/2023.10.18.562872

**Authors:** Olivier Rivoire

## Abstract

The role played by conformational changes in enzyme catalysis is controversial. In addition to examining specific enzymes, studying formal models can help identify the conditions under which conformational changes promote catalysis. Here, we present a model demonstrating how conformational changes can break a generic trade-off due to the conflicting requirements of successive steps in catalytic cycles, namely high specificity for the transition state to accelerate the chemical transformation and low affinity for the products to favor their release. The mechanism by which the trade-off is broken is a transition between conformations with different affinities for the substrate. The role of the effector that induces the transition is played by a substrate “handle”, a part of the substrate that is not chemically transformed but whose interaction with the enzyme is nevertheless essential to rapidly complete the catalytic cycle. A key element of the model is the formalization of the constraints causing the trade-off that the presence of multiple states breaks, which we attribute to the strong chemical similarity between successive reaction states – substrates, transition states and products. For the sake of clarity, we present our model for irreversible one-step unimolecular reactions. In this context, we demonstrate how the different forms that chemical similarities between reaction states can take impose limits on the overall catalytic turnover. We first analyze catalysts without internal degrees of freedom, and then show how two-state catalysts can overcome their limitations. Our results recapitulate previous proposals concerning the role of conformational changes and substrate handles in a formalism that makes explicit the constraints that elicit these features. In addition, our approach establishes links with studies in the field of heterogeneous catalysis, where the same trade-offs are observed and where overcoming them is a well-recognized challenge.

## INTRODUCTION

Two widespread but puzzling features distinguish enzymes from chemical catalysts such as small molecules or large solid surfaces. First, many enzymes undergo conformational changes on the same timescale as their catalytic cycle [1–3], but the role of these conformational changes in catalysis is debated [4–6], notably because our mechanistic understanding of chemical processes suggest that rigid active sites provide optimal environments for chemical transformations [7, 8]. Second, many enzymes catalyze reactions in which the reactant comprises a “handle”, i.e., a non-reactive part that is not transformed chemically but whose interaction with the enzyme is critical to efficient catalytic turnover [9]. Examples include phosphate groups in glycolysis [10], coenzyme A in fatty acid metabolism [11], and amino acid chains extending the cleaved peptide bond in proteolysis [12]. The contribution of these handles is not obvious given Pauling principle [13] that explains catalysis by a specific stabilization of transition states. Since these handles are unchanged, they indeed bind uniformly to substrates and transition states. For multi-molecular reactions, they can accelerate the chemical transformation by bringing and keeping together multiple substrates, but substrate handles are as common in the enzymatic catalysis of unimolecular reactions where this mechanism of catalysis by proximity cannot be invoked [14].

Several unrelated explanations have been proposed to explain conformational changes and substrate handles in enzymatic catalysis. One class of explanations view conformational changes as a means to achieve conflicting geometrical requirements. For instance, a catalytic transformation may be optimized by totally surrounding the reactant with the enzyme, which is incompatible with binding and release [15, 16]. Or, multiple transitions states may be present along the chemical transformation, each requiring a different geometry [17, 18]. Other types of constraints have also been invoked, including demands for substrate specificity, as in the induced-fit model [19, 20], or for regulation, as in many models of allostery [21]. Similarly, several explanations have been proposed for the role of substrate handles. For instance, phosphate groups on many metabolites are justified by the negative charge that they confer, which prevents metabolites to which they are attached from diffusing outside cells [22]. More generally, however, two proposals were made in the 1970s that directly link substrate handles to catalytic efficiency, understood as the rapid completion of a catalytic cycle.

One proposal due to Albery and Knowles arises from their extensive study of triosephosphate isomerase [23], a very efficient enzyme that catalyzes an essential unimolecular reaction in glycolysis, the conversion between two triosephospate isomers that harbor the same phosphate handle. Their explanation invokes chemical and evolutionary constraints and relies on a classification of binding mechanisms contributing to catalysis by the degree of discrimination that they can achieve, with the idea that less discriminative mechanisms are evolutionarily more accessible [24]. From this point of view, the easiest mechanism to evolve is uniform binding to a substrate handle, the term “uniform” referring to the absence of discrimination between the substrate, the transition state and the product. Next, Albery and Knowles considered the possibility of further improvement through “differential binding” where the binding affinity to a transition state is constrained to be intermediate between the binding affinities of the two states preceding and following it. This constraint is motivated by the widespread observation of linear relationships between the affinities to substrates, transition states and products [25]. Finally, they considered improvements through the most general possibility of arbitrary binding to each state, which they called “catalysis of elementary steps”. For triosephosphate isomerase, they argued that uniform binding through the phosphate handle is responsible for most of the improvement over catalysis by a simpler carboxylate base, with differential binding and catalysis of elementary steps making only smaller additional contributions [24]. In their model, a key assumption is that catalysis is present without the handle, and a key variable is the ambient substrate concentration. The increased binding affinity provided by the handle indeed acts to retain the substrate close to the active site until it is chemically transformed, but does not change the activation barrier for the chemical transformation itself. Only for sufficiently low substrate concentrations, when substrate unbinding is limiting, is their model therefore relevant.

In terms of Michaelis-Menten kinetics, uniform binding to the substrate handle increases catalytic efficiency in this first scenario by reducing the Michaelis constant *K*_*M*_ without affecting the catalytic constant *k*_cat_. In many cases, however, altering the interaction of the enzyme with the handle has the very opposite effect: *K*_*M*_ is unchanged but *k*_cat_ is reduced [9]. This puzzling observation motivated Jencks to elaborate a different explanation for the ubiquity of substrate handles. Most relevant to unimolecular reactions is his proposal that the discrimination between a substrate and the transition state can be mainly achieved by destabilizing the substrate rather than by stabilizing the transition state [9, 26]. In this view, the role of the handle is to provide sufficient negative interaction free energy to compensate for the positive free energy involved in substrate destabilization. In Jencks’ words, substrate handles provide a large “intrinsic binding energy” which is not apparent in measured binding energies but is “used as the currency to pay for substrate destabilization” [9].

In contrast to Albery and Knowles’ proposal, Jencks’ proposal is independent of the substrate concentration and involves a lowering of the activation barrier for the chemical transformation. The role of an intrinsic binding energy has been demonstrated in several instances, including triosephosphate isomerase [27, 28]. In particular, Richard and collaborators have shown how the intrinsic energy provided by the interaction with the substrate handle is used to drive many enzymes from a flexible inactive state into a stiff active state through a transition that parallels allosteric transitions: when the substrate in cut in two pieces, the dissociated handle acts as an allosteric effector for the catalysis of the remaining reactive part [29]. In these works, two explanations are given for this mechanism. The first, already mentioned above, is the need to accommodate an open form where the active site is accessible to the substrate, with a closed form where the enzyme optimally encloses it [15, 16]. The second explanation, aligned with the model that we present below, is the need to avoid too tight an association with the substrate [29].

Albery and Knowles’ proposal rests on the quantitative analysis of a model that makes explicit an optimality criterion and the constraints under consideration. In contrast, the conditions under which substrate destabilization is preferable over transition-state stabilization, and the conditions under which an transition between an active and inactive forms are beneficial, have not been formally established. Here, we show how the formalism introduced by Albery and Knowles, known as kinetic barrier diagrams [30], can be extended to account for these mechanisms. Focusing on unimolecular reactions for clarity and because they pose the most significant challenges to explain substrate handles [14], we first derive limits on the cycling time of catalysts that exist only in one conformation. We then show that the capacity of a catalyst to have different affinities for the same ligand when occupying different conformational states, which is one of the hallmarks of allostery [38], can lift some of these limitations. This provides a general formulation of a principle that we previously illustrated with a minimal physics model [39].

Formulating our model with kinetic barrier diagrams has several advantages. Firstly, it offers a rigorous framework that is free from the limitations and inconsistencies of alternative representations based on energy or free energy landscapes [30]. Secondly, it allows us to integrate the explanations of Jencks with those of Albery and Knowles and thus to clarify how they differ but are not exclusive. Finally, kinetic barrier diagrams are not only widely used in studies of enzymes [32–34, 37], but also in studies of non-enzymatic catalysts [35] and in more general studies of non-equilibrium biophysical problems [36]. They therefore provide a common language for biochemists, chemists and biophycists to discuss how enzymes differ from chemical catalysts. In particular, we point out that similar constraints limit the efficiency of single-state catalysts in our model and that of catalytic surfaces in the field of heterogeneous catalysis, where these constraints imply a well-known trade-off known as Sabatier principle [40]. Breaking this trade-off is a well-recognized challenge in this context [41]. Formally demonstrating a mechanism by which this can be achieved is therefore of potential interest beyond the study of enzymes.

## METHODS

### Outline

We first present an informal overview of our results without reference to kinetic barrier diagrams before introducing this framework to provide a more precise presentation.

The main thrust of the model is to expose and analyze the trade-offs that arise between the different steps of a catalytic cycle as a result of the chemical similarity between the reaction states. In the simple context of unimolecular reactions on which we focus, our model distinguishes three reaction states: the substrate *S*, the product *P* and the transition state *S*^*‡*^. Accelerating the chemical transformation requires a catalyst to selectively stabilize the transition state (Pauling principle). In the model, this translates into the requirement that the quantity Δ*G*_*S‡*_ *<* 0 representing the stabilization of the transition state by the catalyst is more negative than the quantity Δ*G*_*S*_ *<* 0 representing the stabilization of the substrate, i.e. Δ*G*_*S‡*_ *<* Δ*G*_*S*_. This condition is necessary, but not sufficient, for a catalytic cycle to be completed faster than a spontaneous reaction. Another important prerequisite is that the product is released sufficiently rapidly. This requires low product stabilization, represented by a sufficiently high value of the quantity Δ*G*_*P*_ *<* 0, representing the stabilization of the product (see Eq. (15)).

The difficulty lies in the fact that the three parameters Δ*G*_*S*_, Δ*G*_*S‡*_ and Δ*G*_*P*_ that define a catalyst in our model cannot generally be modified independently due to the chemical similarity between the three reaction states *S, S*^*‡*^ and *P* : the stabilization of *S*^*‡*^, which is necessary for catalysis, generally implies, at least partly, the stabilization of *S* and *P*, which can be detrimental to the completion of the catalytic cycle. The proposal is that conformational changes provide a means of escaping the trade-offs implied by these constraints.

The model analyzes the consequences of the different forms that these chemical constraints can take, formalized by different types of correlation between the values of the three parameters Δ*G*_*S*_, Δ*G*_*S‡*_ and Δ*G*_*P*_. The simplest case assumes that these three parameters are constrained to have the same value, Δ*G*_*S*_ = Δ*G*_*S‡*_ = Δ*G*_*P*_. Such a uniform binding cannot ensure catalysis per se, since the transition state is not stabilized with respect to the substrate. Albery and Knowles noted, however, that the addition of uniform binding to a pre-existing catalytic mechanism can be beneficial [24]. Formally, this amounts to assuming a pre-existing catalytic mechanism defined by certain 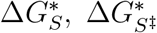 and 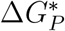 values, and then examining how adding a common Δ*G*_*u*_ *<* 0 value to these three parameters can contribute to faster completion of a catalytic cycle. This is shown to be the case if the substrate concentration is sufficiently low. The addition of this uniform binding energy Δ*G*_*u*_ is the justification for substrate handles proposed by Albery and Knowles.

Another form of chemical constraint is to assume that Δ*G*_*S‡*_ is constrained to be intermediate between Δ*G*_*S*_ and Δ*G*_*P*_. This form of differential binding reflects the widespread observation of linear free-energy relationships in chemistry [25]. It is formalized by taking Δ*G*_*S*_ and Δ*G*_*P*_ as independent parameters and assuming that Δ*G*_*S‡*_ takes an intermediate value between the two: Δ*G*_*S‡*_ = (1*− λ*)Δ*G*_*S*_ + *λ*Δ*G*_*P*_ where 0 *< λ <* 1 represents the degree of similarity of the transition state to the product. We show that in this case Δ*G*_*S*_ and Δ*G*_*P*_ can be chosen to achieve catalysis, but that the activation barrier cannot be reduced by a factor greater than two – or more precisely, by a factor greater than 1+*λ* (see Eq. (21)). This excludes, in particular, diffusion-limited catalysis where activation barriers are completely annihilated.

To overcome this limitation, it is necessary to decouple the two catalytic steps in trade-off, namely the chemical transformation *CS → CP* and the release of the product *CP → C* + *P*. One mechanism for achieving this is to have the catalyst adopt two different states with different binding properties during the catalytic cycle, a state *C*_1_ for the chemical transformation *C*_1_*S → C*_1_*P* and a state *C*_0_ for the release of the product *C*_0_*P → C*_0_ + *P*. Such a two-state catalyst is described in our model by 7 parameters: the stabilization of the reaction states in each state of the catalyst, 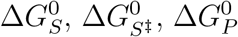, and 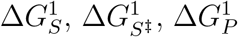, and the difference of free energy between the two states of the catalyst in the absence of any reactant, Δ*G*_*C*_. Although these 7 parameters expand the range of possible catalysts, we show that when each catalyst state is subject to differential binding constraints, catalysis is still limited to lowering the activation barriers by a factor of at most two.

We propose, however, that another type of correlation between the stabilization of reaction states better reflects the constraints to which enzymes are subjected. The idea is that the transition state can be stabilized more strongly than the substrate and product, but only at the expense of a greater stabilization of these two states. This describes, for example, a situation where effective stabilization of the transition state requires precise, fixed positioning of the substrate, which can also only be achieved by strong binding of both the substrate and the product. Or a situation of strain catalysis where high strain is only possible at the cost of a tight binding of both the substrate and the product. We formalize this type of constraint, which we call discriminative binding, by assuming that in any given catalyst state, Δ*G*_*S*_, Δ*G*_*S‡*_ and Δ*G*_*P*_ are constrained to satisfy Δ*G*_*S*_ = Δ*G*_*P*_ = Δ*G*_*u*_ and Δ*G*_*S‡*_ = (1 + *α*)Δ*G*_*S*_ where *α* controls the increment in the stabilization of the transition state achieved by stabilizing the substrate and product. For example, we previously presented a simple model of strain catalysis where *α* = 1 [39].

Under this form of chemical constraint, which we call discriminative binding, we show that a single-state catalyst can at most lower the activation energy by a factor of 1 + *α* (see Eq. (24)). However, a two-state catalyst can overcome this limitation and eventually reduce the activation barrier altogether. This is achieved through a particular design in which one state is inactive with 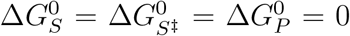 while the other strongly binds the transition state. The two remaining parameters 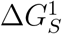 and Δ*G*_*C*_ must be well chosen for this mechanism to be effective: we must have 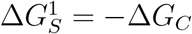 so that the energy cost of the conformational change is offset by the substrate’s affinity for the catalytically active state, and Δ*G*_*C*_ must exceed the activation energy to abolish it (see Eq. (36)).

In this model, 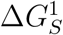therefore plays the role of the intrinsic binding energy of Jenck’s proposal. It is not directly observed as binding free energy, as it is compensated for by Δ*G*_*C*_, but is essential for stabilizing the transition state. Our model, however, differs from a mechanism of pure substrate destabilization, which we find to have limited catalytic efficiency (Supplementary Material (SM) Sec. 6). Our model, on the other hand, is analogous to an allosteric mechanism where a substrate handle provides the intrinsic binding energy 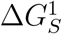 to activate the enzyme through a transition to a different conformation of higher free energy Δ*G*_*C*_.

One of the merits of our model is to establish a direct link with observations in heterogeneous catalysis involving solid (single-state) surfaces. The trade-off between chemical transformation and product release is well known in this context to be a central limitation to the efficient completion of catalytic cycles. It is usually represented by Volcano diagrams showing that catalytic efficiency exhibits a non-trivial optimum as a function of the catalyst’s affinity for the substrate [40]. This optimum follows Sabatier principle: the catalyst must bind sufficiently strongly to the transition state, but not too strongly. Our model recapitulates these observations when considering a single catalyst subjected to either differential or discriminative binding constraints. Studies on heterogeneous catalysts have in fact gone further in characterizing the nature of the chemical constraints at play, by showing scaling relationships [42], which can take exactly the form of the discriminative binding constraints that we propose (see SM Sec 7).

### Model

#### Spontaneous reaction

We consider for simplicity a unimolecular reaction described by a single-step mechanism,

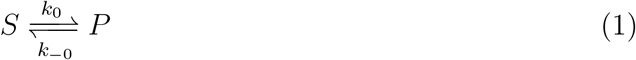

where *S* represents the substrate, *P* the product, *k*_0_ the first-order rate constant for the forward reaction and *k*_*−*0_ the first-order rate constant for the reverse reaction. The rate of product formation is then *v*_0_ = *∂*[*P*]*/∂t* = − *∂*[*S*]*/∂t* = *k*_0_[*S*] − *k*_*−*0_[*P*] where [*S*] and [*P*] are, respectively, the concentrations of substrate *S* and product *P*. To model a cellular context, we study this reaction in a non-equilibrium steady state where these concentrations are maintained at fixed values.

To reason about catalysis, it is convenient to consider (free) energies rather than rates [35]. We therefore introduce a parametrization of the two rates *k*_*±*0_ by two free energies, an activation free energy 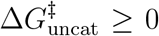 and a free energy of formation of one molecule 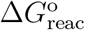such that

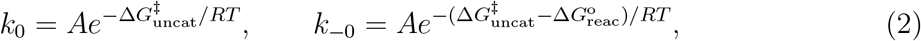

where *R* is the universal gas constant, *T* the temperature, and *A* a frequency factor (*A* = *k*_*B*_*T/h* in transition state theory, *k*_*B*_ being Boltzmann constant and *h* Planck constant). To simplify the formulas, we set the unit of energy to have *RT* = 1 and the unit of time to have *A* = 1. The two quantities 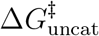and 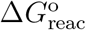are, by definition, independent of the concentrations of *S* and *P*. 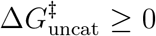represents a positive activation energy for the forward reaction *S → P* while 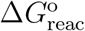is a free energy of formation related to the equilibrium constant *K*_eq_ = *k*_0_/*k*_−0_ by 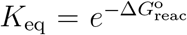 (note that we define 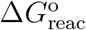*per molecule* rather than *per mole* as more common in chemistry). It is also convenient to introduce the free energy of reaction 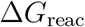when the substrate and product concentrations are fixed to the arbitrary values [*S*] and [*P*],

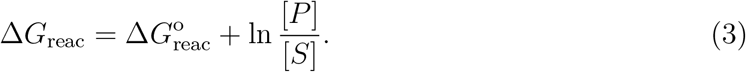

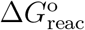can be of any sign, and we only need to impose 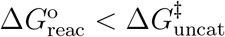for the reverse reaction *P → S* to have a positive activation energy, and therefore for the two states *S* and *P* to be well defined. These parameters for the spontaneous reaction are represented in a kinetic barrier diagram [30, 34] with three states, the two stable states *S* and *P*, whose levels differ by 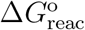, and a transition state *S*^*‡*^ whose level differs from that of *S* by 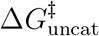(Fig. 1A).

**FIG. 1:**
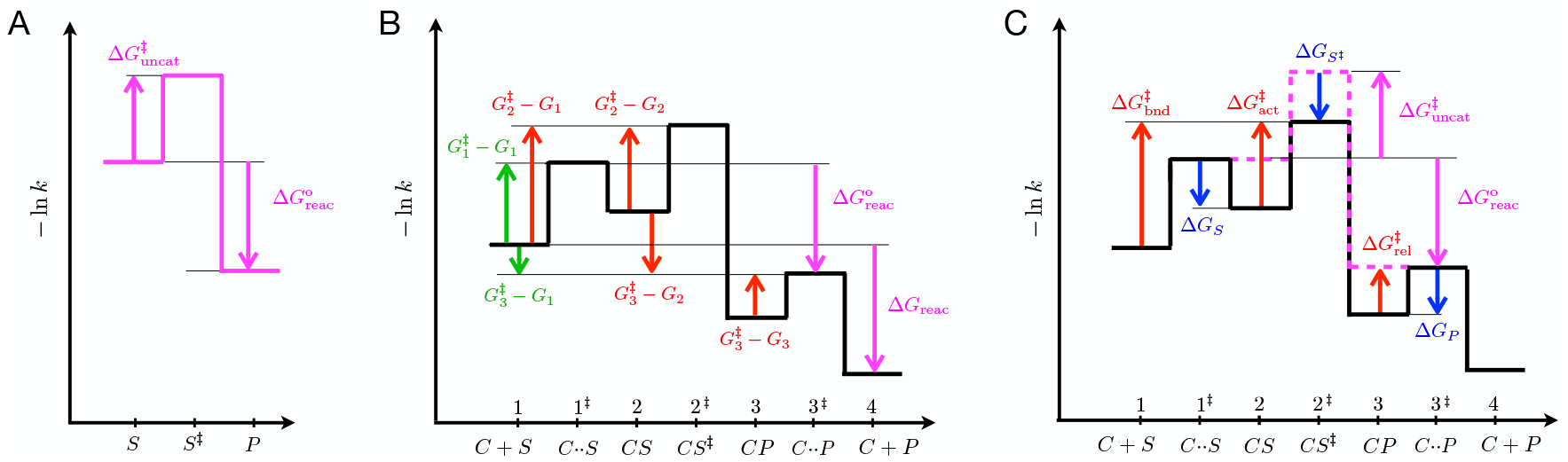
Kinetic barrier diagrams. **A**. Diagram for the spontaneous reaction *S ⇌ P*, described by two stable states, *S* and *P*, a transition state, *S*^*‡*^, and two parameters, an activation barrier and a reaction barrier 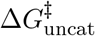and a reaction barrier 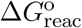. Here 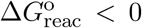but 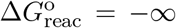 if considering an irreversible reaction (more generally, 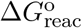can be of any sign as long as 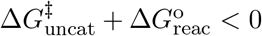). **B**. Diagram for the catalytic process described by Eq. (4). The stable states, *C* + *S, CS, CP* and *C* + *P* are represented as local minima with energies *G*_*i*_ (*i* = 1, 2, 3, 4), separated by transition states *C··S, CS*^*‡*^ and *C··P*, with energies 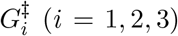. The heights of the barriers represent the transition rates. For instance, the rate from *CS* to *CP* 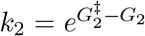 while the reverse rate from *CP* to *CS* is 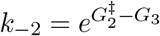. The cycling time *T*_*c*_ expressed in Eq. (9) depends on the forward barriers between successive states, 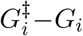, as well as on the forward barriers between non-successive states 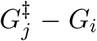 with *j > i*. In total, this corresponds to the 6 barriers represented by green or red arrows. Of these 6 barriers, 2 are set by extrinsic parameters independent of the catalyst (in green) and 4 are modulated by parameters intrinsic to the catalyst (in red). Some barriers may have negative values (downward-pointing arrows) and therefore not constitute barriers *stricto sensu*. In particular, 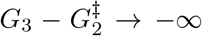 when the reaction is irreversible. Note also that *G*_4_ *→ −∞* when products are maintained at vanishing concentration. **C**. When considering irreversible reactions, only 3 barriers are dependent on properties of the catalyst, 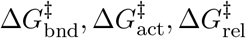, represented by the 3 red arrows. We describe the properties of the catalyst by three intrinsic parameters, Δ*G*_*S*_, Δ*GS*_*‡*_, Δ*G*_*P*_, represented by the 3 blue arrows. They are defined by making a comparison with a non-interacting catalyst which differs from an interacting catalyst in the internal section of the diagram, where its profile is that of the spontaneous reaction (pink dotted lines).

For clarity, we make two further simplifying assumptions: no product is present, [*P*] = 0, and the reaction is irreversible, *k*_*−*0_ = 0 or, equivalently, 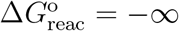 [a generalization to arbitrary 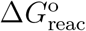 is presented in the Supplementary Material (SM)]. These assumptions imply that the rate of product formation due to the spontaneous reaction is simply *v*_0_ = *k*_0_[*S*].

#### Catalysis

Catalysis occurs if a substrate is converted more quickly in the presence than in the absence of a substance – the catalyst – which is left unchanged in the process. We first consider a catalyst *C* with no internal degree of freedom that follows a catalytic cycle with two intermediate states, described by a Markov chain of the form

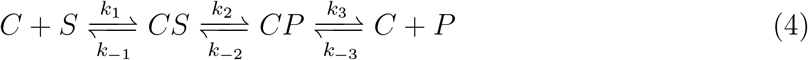

where *k*_1_ = *k*_*D*_[*S*] and *k*_*−*3_ = *k*_*D*_[*P*] are pseudo-first-order rate constants that depend on the ambient concentrations of substrate and product and on a diffusion rate constant *k*_*D*_, while the other rates *k*_*±i*_ are first-order rate constants that depend on properties of the catalyst.

Part of the confusion surrounding the role of conformational changes in catalysis stems from the different definitions of catalysis [6]. This definition is sometimes limited to the chemical transformation from *CS* to *CP*. As the selective pressures exerted on enzymes act on the entire catalytic cycle, we consider here a measure of catalytic efficiency that takes into account each step of the Eq. (4). To do this, we quantify catalytic efficiency by the average time *T*_*c*_ to complete a full catalytic cycle, i.e. to reach *C* + *P* from *C* + *S* in Eq. (4). The smaller the cycling time *T*_*c*_, the more efficient catalysis. In conditions where [*P*] = 0, this cycling time is equivalent to the catalytic efficiency *y* introduced by Albery and Knowles [43]. If, furthermore, the contribution of the spontaneous reaction to the rate of product formation, *v* = *∂*[*P*]*/∂t*, is negligible, *T*_*c*_ is equivalent to [*C*]*/v*, where [*C*] is the total concentration of free and bound catalysts [44, 45]. For the catalytic cycle described by Eq. (4), the rate of product formation follows the Michaelis-Menten equation, *v* = *k*_cat_[*S*][*C*]*/*(*K*_*M*_ + [*S*]) [46], and we can therefore express the dependence of *T*_*c*_ on the substrate concentration [*S*] in terms of a catalytic constant *k*_cat_ and a Michaelis constant *K*_*M*_, as

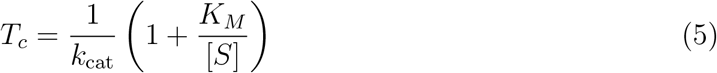

where, for the catalytic cycle described by Eq. (4) (see SM Sec. 1 or [24]),

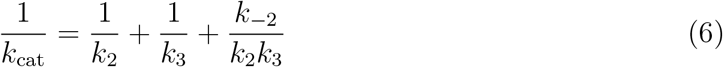

and

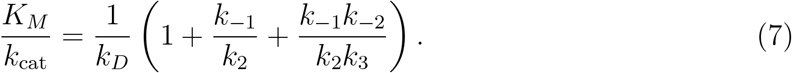

We assumed here *N* = 2 intermediate states, *CS* and *CP*, but Eq. (5) extends to unidimensional chains of transitions with an arbitrary number *N* of intermediate states, with appropriate redefinitions of *k*_cat_ and *K*_*M*_ (see SM Sec. 1).

Since *T*_*c*_ quantifies the time to complete a catalytic cycle with no reference to the spontaneous reaction, its value does not reveal if catalysis is taking place, i.e., if the reaction in the presence of the catalyst is faster than in its absence. In particular, as *T*_*c*_ represents a turn-over time *per catalyst*, it is *not* comparable to the mean spontaneous reaction time 1*/k*_0_ *per substrate*. To assess the presence of catalysis, we must either compare the reaction time 1*/k*_0_ *per substrate* in the absence of catalysts to another reaction time in the presence of catalysts, or compare the cycling time *T*_*c*_ *per catalyst* in the presence of the substrate of interest to a another cycling time where the catalyst is substituted for an inactive substance. The two approaches lead to the same simple criterion valid for any number *N* of intermediate states: catalysis occurs if and only if *k*_0_ *< k*_cat_, where *k*_cat_ is the catalytic constant in Eq. (6). Remarkably, this criterion is independent of *K*_*M*_, whose value impacts the cycling time *T*_*c*_ but has no bearing on the occurence of catalysis *per se* [45].

As for the spontaneous reaction, we can re-parametrize the elementary rates *k*_*±i*_ in Eq. (4) with free energies and represent the catalytic process in a kinetic energy diagram. To this end, each transition is associated with a transition state. The first transition 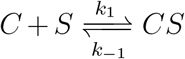 is associated with a first transition state (*i* = 1) denoted *C··S* to represent a substrate just about to bind to the catalyst. The second transition 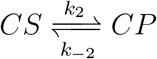 is associated with a second transition state (*i* = 2) denoted *CS*^*‡*^ to represent the transition-state-catalyst complex. The third transition 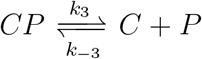, finally, is associated with a third transition state (*i* = 3) denoted *C··P* to represent a product just about to be released from the catalyst. We define free energies *G*_*i*_ for the stable states (*i* = 1 for *C* + *S, i* = 2 for *CS, i* = 3 for *CP* and *i* = 4 for *C* + *P*) and 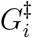for the transition states (*i* = 1, 2, 3), so that

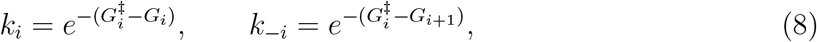

which allows the catalytic cycle of Eq. (4) to be represented by a kinetic barrier diagram (Fig. 1B) [30, 34]. In this diagram, the energy difference between the last state *C* + *P* and the first state *C* + *S* coincides with the free energy change Δ*G*_reac_ defined in Eq. (3) (due to the assumption that the substrate and the product have same diffusion constant *k*_*D*_).

In terms of these free energies, the cycling time *T*_*c*_ takes, when assuming [*P*] = 0, a simple form (see SM Sec. 1 or [35]),

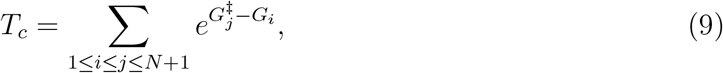

where the sum is over each pair *i≤ j* of a transition state *j* following a stable state *i*. Typically, one term dominates the sum and 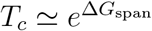where 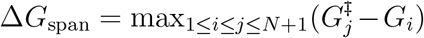 is known as the energetic span [35]. We shall work in this approximation where estimating the cycling time amounts to estimating the limiting barrier 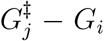that determines the energetic span Δ*G*_span_. Importantly, this limiting barrier is not necessarily associated with a limiting step (*i* = *j*) but can involve a transition state that does not follow immediately the intermediate state (*i < j*). When *N* = 2, (*N* + 1)(*N* + 2)*/*2 = 6 barriers, represented by the green and red vertical arrows in Fig. 1B, have to be compared to determine which is largest. Some of these barriers, however, may have negative values and be therefore negligible. When considering an irreversible reaction, or more generally when considering as in Fig. 1B a reaction with a large activation barrier for the reverse reaction, this is the case of the two barriers 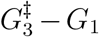and 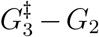, represented in Fig. 1B by downward-pointing arrows.

Similarly to *T*_*c*_, the catalytic constant *k*_cat_ that defines whether catalysis is present (if *k*_cat_ *> k*_0_) is given by

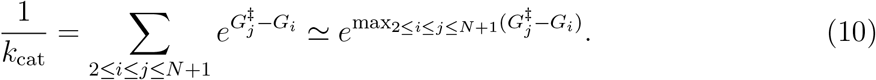

With *N* = 2 intermediate states, *k*_cat_ is therefore determined by the largest of *N* (*N* +1)*/*2 = 3 barriers.

#### Intrinsic and extrinsic barriers

When considering constraints on catalytic efficiency, an important distinction is between intrinsic barriers which depend on properties of the catalyst (in red in Fig. 1), and extrinsic barriers which do not (in green in Fig. 1), and depend instead exclusively on the parameters 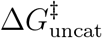and 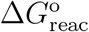of the spontaneous reaction and on the ambient concentrations [*S*] and [*P*]. In the catalysis of an irreversible reaction with no product and *N* = 2 intermediates, only three barriers are intrinsic and non-negative, represented by the three red upward-pointing arrows in Fig. 1B. Given the essential role of these barriers in what follows, it is convenient to give them short names (Fig. 1C),

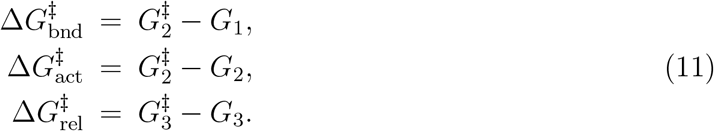

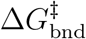is a barrier involving both binding and activation, associated with the transition *C* + *S → CP*, and is all the higher that the substrate concentration is lower (small *G*_1_) and the activation energy is higher (large *G*_2_). Δ*G*_act_ is an activation barrier for the chemical transformation in presence of the catalyst, controlling the transition *CS → CP*. 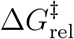, finally, is associated with product release, and controls the transition *CP → C* + *P* .

With these notations, Eq. (9) can be rewritten as

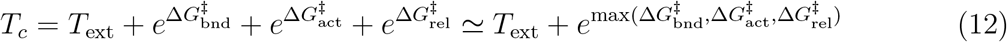

where *T*_ext_ a lower bound on the cycling time that is set by the extrinsic parameters and is therefore independent of the catalyst itself; for irreversible reactions, *T*_ext_ is simply the mean time needed for a substrate to diffuse towards a catalyst. Similarly,

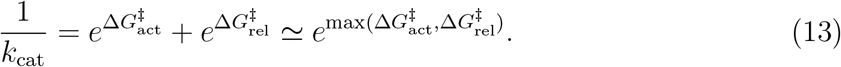

#### Intrinsic parameters

If the three intrinsic barriers 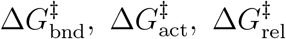can be lowered arbitrarily, perfect catalysis with a minimal cycling time *T*_*c*_ = *T*_ext_ is achievable. The difficulties for a catalyst to discriminate between the reaction states *S, S*^*‡*^ and *P* (Fig. 1A) may, however, prevent this optimum to be reached. To analyze the trade-offs at play, we need to relate the three intrinsic barriers 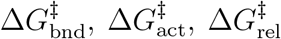 to physical parameters reporting the affinity of the catalyst to the three reaction states.

To this end, we take as reference a non-interacting catalyst subject to the same extrinsic conditions. By definition, its kinetic barrier diagram differs only in its internal section, as represented by the pink dotted lines in Fig. 1C: it has an activation barrier identical to that of the spontaneous reaction 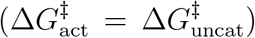, no barrier for release 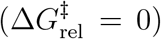 and the binding/activation barrier is entirely controlled by diffusion 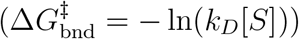. An actual catalyst differs from this non-interacting catalyst by the extent to which the free energies of the three states *CS, CS*^*‡*^ and *CP* are lowered, which we quantify with the three intrinsic parameters Δ*G*_*S*_, Δ*G*_*S‡*_ and Δ*G*_*P*_ represented by blue arrows in Fig. 1C. These three parameters, which can be thought as binding free energies are, by definition, independent of reactant concentrations and have necessarily negative values.

In terms of the three intrinsic parameters Δ*G*_*S*_, Δ*G*_*S‡*_, Δ*G*_*P*_, the three intrinsic barriers controlling *T*_*c*_ are given by

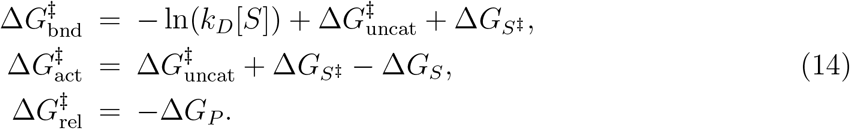

We use below these expressions to study how the limiting barrier max 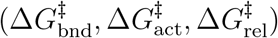 in Eq. (12) is minimized as Δ*G*_*S*_, Δ*G*_*S‡*_ and Δ*G*_*P*_ are varied.

#### Conditions and fundamental limits to catalysis

From Eqs. (2) and (13), it follows that catalysis (*k*_cat_ *> k*_0_) requires max 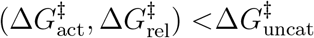, which, given Eq. (14), corresponds to

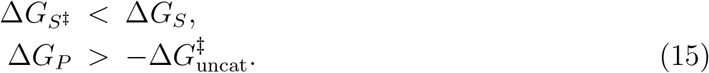

The first condition embodies Pauling principle [47]: the catalyst must bind more strongly to the transition state than to the substrate to reduce the activation energy. The second condition imposes the product not to bind too strongly, to allow for efficient product release. Neither minimizing each of the three barriers in Eq. (14) nor satisfying Eq. (15) involve any trade-off: minimizing Δ*G*_*S‡*_ while maximizing Δ*G*_*S*_ and Δ*G*_*P*_ contributes to minimize each barrier in Eq. (14) and permits to satisfy Eq. (15). In this model, the maximal value of *k*_cat_ (*k*_cat_ = 2) is for instance achieved with Δ*G*_*S*_ = 0, 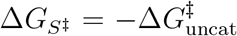, Δ*G*_*P*_ = 0. This limit corresponds to so-called perfect catalysis, where the limiting process is the diffusion of a substrate towards a catalyst [43].

## RESULTS

### Constraints and limitations on single-state catalysis

We propose to understand general design principles of enzymes as arising from generic but non-thermodynamical constraints to which the parameters Δ*G*_*S*_, Δ*G*_*S‡*_ and Δ*G*_*P*_ are subject. We ignore constraints from geometry, specificity or regulation, and focus instead on constraints arising from the chemical similarity of the three reaction states *S, S*^*‡*^ and *P*. We model these constraints by imposing a positive correlation between Δ*G*_*S*_, Δ*G*_*S‡*_ and Δ*G*_*P*_. First, we follow Albery and Knowles and re-analyze the cases of uniform binding, where the three free energies are imposed to be the same, and of differential binding, where Δ*G*_*S‡*_ is assumed to lie between Δ*G*_*S*_ and Δ*G*_*P*_ [24]. Next, we introduce and justify a new type of constraint that we call discriminative binding, where the specificity to the transition state Δ*G*_*S‡*_*−* Δ*G*_*S*_ is positively correlated to the affinities to the substrate and product Δ*G*_*S*_ and Δ*G*_*P*_. Two other constraints are also analyzed in Supplementary Material, one capturing the notion of substrate destabilization proposed for enzyme catalysis [26] and another capturing the scaling laws observed in heterogeneous catalysis [42, 48]. Throughout this section, we assume single-state catalysts described by Eq. (4) before analyzing in the next section the benefit of catalysts with an internal degree of freedom.

#### Single-state uniform binding

The most restrictive constraint is to assume uniform binding, where the interaction between the reactant and the catalyst is independent of the state of the reactant and described by a single parameter Δ*G*_*u*_ *≤* 0, such that

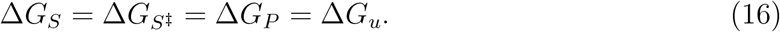

This constraint represents, in particular, the interaction of an enzyme with a non-reactive substrate handle, which is independent of the chemical state of the reactive part of the substrate. Since catalysis (*k*_cat_ *> k*_0_) requires 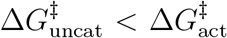 and since uniform binding leaves 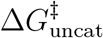unchanged [see Eq. (14)], such uniform binding cannot confer catalysis [14]. As proposed by Albery and Knowles [24], it can, however, be beneficial when complementing a pre-existing catalytic mechanism. Adding uniform binding Δ*G*_*u*_ *≤* 0 to a pre-existing catalytic mechanisms with intrinsic barriers 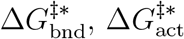 and 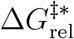 indeed leads to

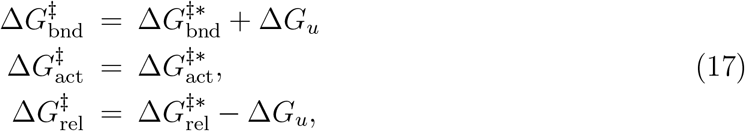

i.e., a reduction of the binding/activation barrier 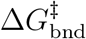 at the expense of an equal increase of the release barrier 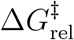. This is advantageous when 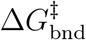 is limiting. Since the lower the [*S*] substrate concentration, the higher 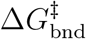, this scenario depends critically on the substrate concentration and applies when this concentration is sufficiently low, namely when 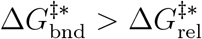 (Fig. 2 and SM Sec. 4). The optimal value of Δ*G*_*u*_ is reached when 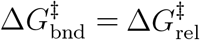. Albery and Knowles argued that this effect explains most of the improvement of triosephosphate isomerase provides over a non-enzymatic catalyst [24].

**FIG. 2:**
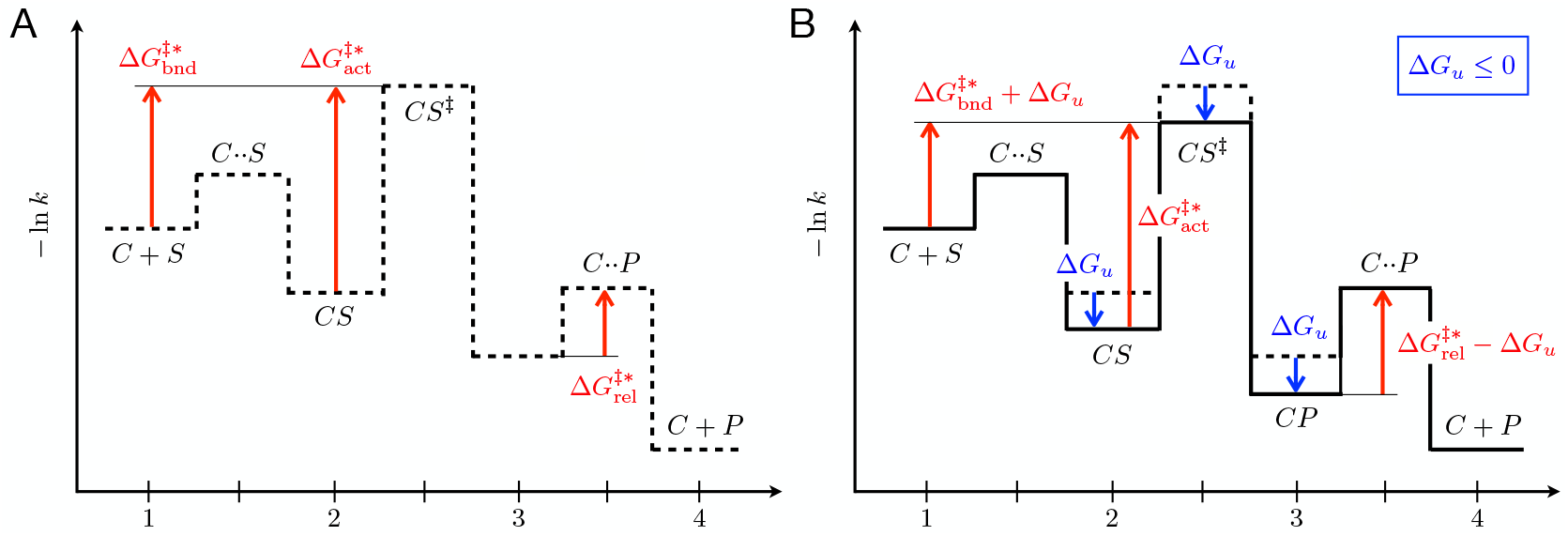
Single-state uniform binding. **A**. A pre-existing catalytic mechanism is assumed where 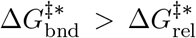. **B**. Adding uniform binding to this pre-existing mechanism lowers 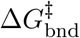 at the expense of a larger 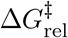. Given 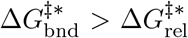, the value of Δ*G*_*u*_ *<* 0 that minimizes the maximum of these two barriers is such that 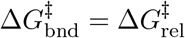. This leads to a shorter cycling time, but 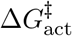is unchanged, and, as in the case represented here, may remain the limiting barrier.

#### Single-state differential binding

A less stringent constraint than uniform binding is differential binding which accounts for an empirical observation known in chemistry as the Bell-Evans-Polanyi principle [25, 49]. This principle generally relates the difference of activation energies of two related reactions, 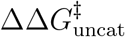, to the difference of their reaction energies, 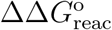, by a linear relationship 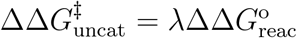 with 0 *≤ λ ≤* 1. In our model where we are comparing reactions in the context of different catalysts, this amounts to assuming that Δ*G*_*S‡*_ is constrained to lie between Δ*G*_*S*_ and Δ*G*_*P*_, which can also be expressed by a linear relationship,

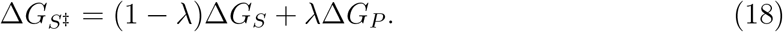

This constraint formalizes the notion that the transition state *S*^*‡*^ has chemical properties that are intermediate between those of the substrate *S* and the product *P*. In this view, *λ* reports the degree to which the transition state *S*^*‡*^ is more similar to the product *P* than to the substrate *S*. Two independent intrinsic parameters are left, Δ*G*_*S*_ and Δ*G*_*P*_ .

In contrast to uniform binding, differential binding can confer catalysis on its own, but, as we now show, only to a limited extent. To derive this limitation, we express the two kinetic barriers that control *k*_cat_ as a function of the two tunable parameters Δ*G*_*S*_ and Δ*G*_*P*_,

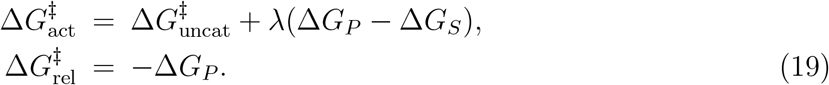

This makes apparent a trade-off between activation and release, which depend with opposite signs on Δ*G*_*P*_. Increasing |Δ*G*_*P*_ | decreases 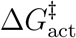 but increases 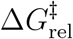 (since Δ*G*_*P*_ *≤* 0). This trade-off reflects a well-known principle in heterogeneous catalysis, Sabatier principle, which states that an optimal catalyst must strike a balance between sufficient strong interaction to activate the reactant and sufficient low interaction to facilitate product release [50, 51].

If |Δ*G*_*P*_ | is low, the barrier limiting *k*_cat_ is 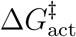 while if it is large it is 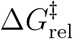. The maximal value of *k*_cat_ is obtained when the two barriers 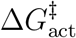and 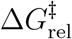are equivalent, which corresponds to

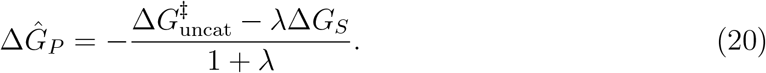

Given Δ*G*_*S*_ *≤* 0, this implies an upper bound on *k*_cat_,

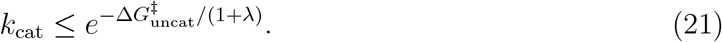

Under constraints of differential binding, catalysis can thus reduce the activation barrier 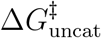 by a factor (1 + *λ*) *≤* 2 at most, which excludes in particular perfect catalysis. This conclusion is verified numerically when sampling the space of possible parameters (SM, Fig. S3A).

#### Single-state discriminative binding

Here we introduce another form of constraint between Δ*G*_*S*_, Δ*G*_*S‡*_ and Δ*G*_*P*_, which we propose to better capture an essential trade-off to which enzymes are subject. Perhaps the simplest mechanism by which binding can contribute to enzymatic catalysis is indeed a precise and rigid positioning of the reactant, in a configuration that defines an optimal chemical environment for the reaction. However, such precise positioning typically necessitates tight binding of the substrate (high |Δ*G*_*S*_|), which cannot be achieved through interactions limited to the small reactive part of the substrate. Instead, it must involve other, non-reactive parts of the substrate that are also present in the product, implying that |Δ*G*_*P*_| is also high. This type of catalytic mechanism therefore involves a trade-off between the specificity Δ*G*_*S‡*_*−* Δ*G*_*S*_ and the affinities Δ*G*_*S*_ and Δ*G*_*P*_. A similar trade-off is expected if alternatively considering catalysis through a strain mechanism, where again high strain is typically coupled to tight binding, irrespectively of the reaction state. To formalize in simple terms this trade-off, we propose to consider that Δ*G*_*S*_, Δ*G*_*S‡*_ and Δ*G*_*P*_ are dependent on a single degree of freedom Δ*G*_*u*_ *≤* 0 with

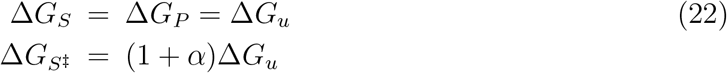

where *α ≤* 0 is a fixed parameter that quantifies the potential for transition-state specificity, with uniform binding (no specificity) corresponding to the limit *α →* 0. Here, Δ*G*_*u*_ represents uniform binding to the substrate and product, but not to the transition state for which the additional contribution *α*Δ*G*_*u*_ is present. We previously studied a simple physics model which displays this type of constraints with *α* = 1 [39]. More generally, we could assume Δ*G*_*S‡*_ = Δ*G*_*u*_ + *f* (Δ*G*_*u*_) where *f* (Δ*G*_*u*_) *≤* 0 is an increasing function of Δ*G*_*u*_ that can take arbitrary low values. However, as in the case of differential binding where we limited the analysis to a linear relationship, the phenomenology is already captured by the linear function *f* (Δ*G*_*u*_) = *α*Δ*G*_*u*_.

Under the constraints of Eq. (22), which we call discriminative binding, the two barriers controlling *k*_cat_ are

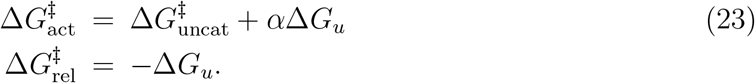

A trade-off consistent with Sabatier principle is again obtained, where a decrease of the activation barrier is coupled to an increase of the release barrier. As previously, the minimum of max 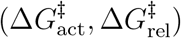is obtained when 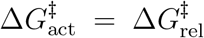which corresponds to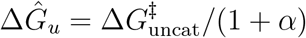. This implies an upper bound on *k*_cat_, namely

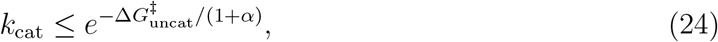

and therefore a lower bound on the cycling time, as can also be verified numerically (SM, Fig. S3C). In particular, perfect catalysis is again excluded under this scenario.

### Constraints and limitations on two-state catalysis

Enzymes can adopt different conformations with different binding free energies for the same ligand, a property that is key to allostery [38]. Here, we analyze how the presence of two such conformations can contribute to overcome the limitations of catalysts with a single conformation. We take the two states of the catalyst, denoted *C*_0_ and *C*_1_, to be associated with different sets of binding free energies, respectively 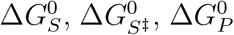 and 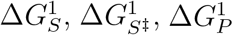, and we assume that constraints due to chemical similarity between reaction states apply independently in each state of the catalyst. *C*_0_ is taken to represent the state of lowest free energy and we describe the transition between the two catalytic states similarly to the spontaneous reaction as

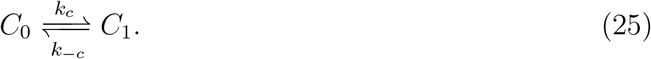

Again, we parametrize the rates with a free energy difference Δ*G*_*C*_ *≥* 0 and an internal barrier 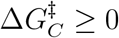, such that (Fig. 3A)

**FIG. 3:**
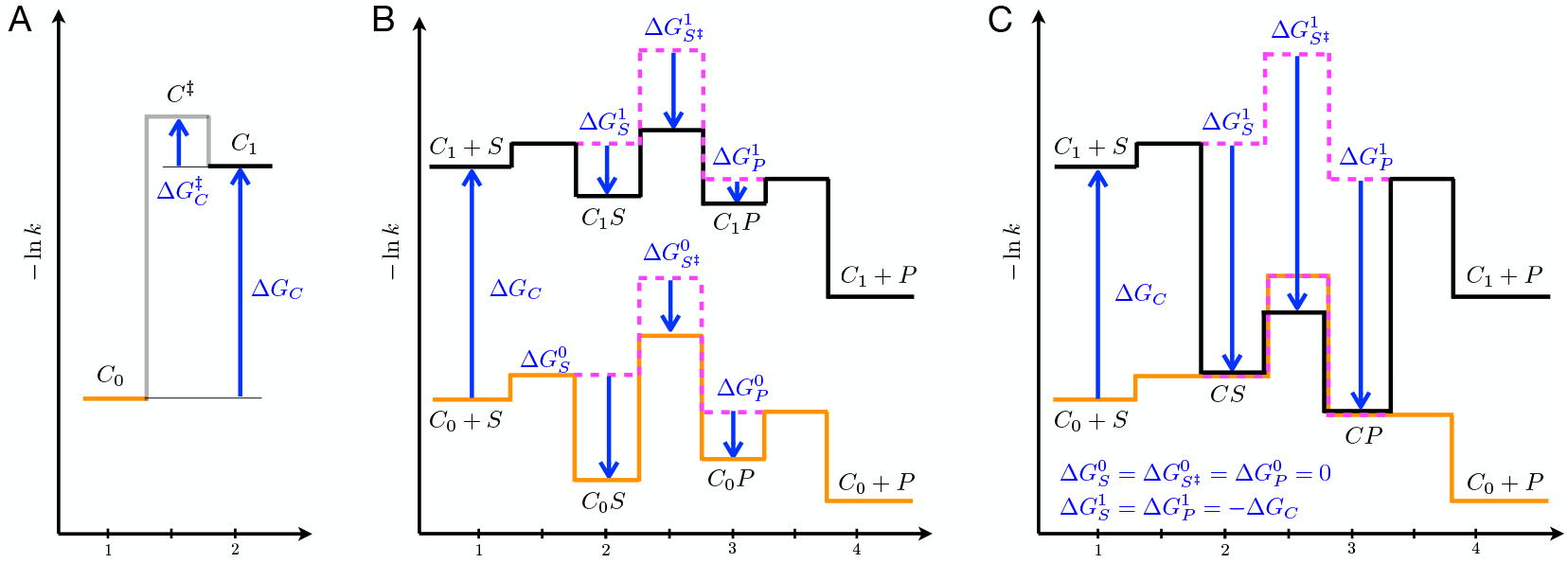
Two-state catalysis. **A**. A catalyst can be in two states, a low-free-energy conformation *C*_0_ and a high-free-energy conformation *C*_1_. The transitions between these states are parametrized by the free energy differences Δ*G*_*C*_ *≥* 0 and 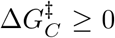. **B**. Kinetic barrier diagram representing the transitions within each state of a two-state catalyst. The transitions between the two conformations – corresponding to the horizontal transitions of the two-dimensional network of Eq. (27) – are not represented, which would require introducing a third dimension. As in Fig. 1, the intrinsic parameters are represented by blue arrows and the energy levels for a non-interacting catalyst subject to the same extrinsic conditions are represented by pink dotted lines. **C**. Particular case where *C*_0_ is inactive 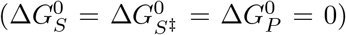 and where binding in *C*_1_ compensates for the cost of the conformational change 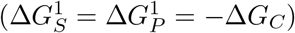, so that *C*_0_*S* and *C*_1_*S* have same free energy, and so do *C*_0_*P* and *C*_1_*P*. Assuming further that 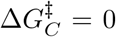, each of these pairs of states can be treated as a single state, here denoted *CS* and *CP* .

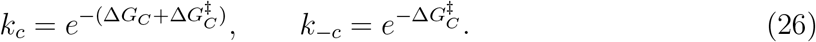

Generalizing Eq. (4), a catalyst in presence of substrates can be in eight possible states that are interconnected in a two-dimensional network of transitions of the form

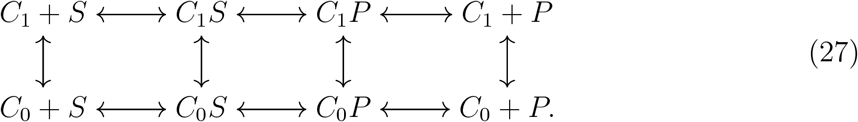

The number of intrinsic parameters, which was 3 for single-state catalysts, is 8 for two-state catalysts, namely 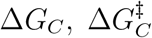and 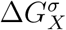for *X* = *S, S*^*‡*^, *P* and *σ* = 0, 1 (blue arrows in Fig. 3A-B).

The derivation of an analytical formula for the cycling time in the most general case is laborious, but to demonstrate the possibility of reaching the diffusion limit, it suffices to expose a particular case where this limit is reached. This particular case can be obtained under conditions justifying approximations that simplify the analysis. First, the network of Eq. (27) contains many paths from *C*_0_ +*S* to *C*_0_ +*P* but one typically drives most of the flux, which makes possible an approximation of the dynamics by a one-dimensional succession of transitions. This is the case in the limit in which we focus here, where Δ*G*_*C*_ is sufficiently large for the states *C*_1_ + *S* and *C*_1_ + *P* to have negligible probabilities compared to *C*_0_ + *S* and *C*_0_ + *P*, and where 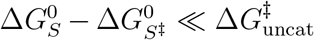 so that *C*_*0*_ is catalytically inactive and the transition *C*_0_*S → C*_0_*P* therefore negligible. In this limit, the dominant path in Eq. (27) is

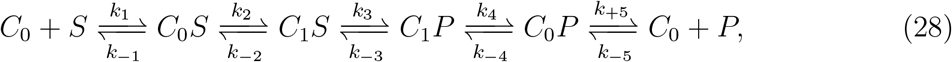

and the cycling time can be computed using Eq. (9) with *N* = 4 intermediate states.

With further assumptions, however, the dynamics can be described by an even simpler model with just *N* = 2 intermediate states. For *C*_0_ + *S → C*_1_ + *S* to be negligible but not *C*_0_*S → C*_1_*S*, the “cost” Δ*G*_*C*_ of the conformational change must indeed be offset by a nearly equivalent gain in binding free energy, with 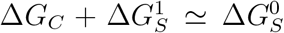. When this compensation takes place and when 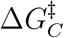 is negligible, the interconversion *C*_0_*S* ⇌ *C*_1_*S* occurs on a fast time scale, and the two states *C*_0_*S* and *C*_1_*S* can be treated as a single state *CS*. If, further, 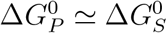 and 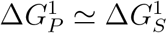, as it is necessarily the case when considering either uniform or discriminative binding, the same argument applies to the interconversion *C*_0_*P ⇌ C*_1_*P*, and the number of intermediate states is reduced to *N* = 2. Under these different assumptions that may be summarized by

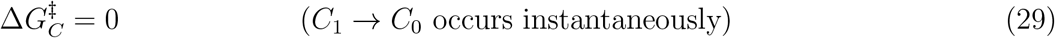

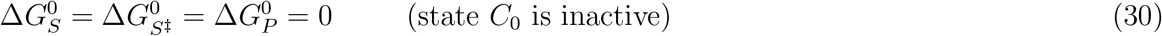

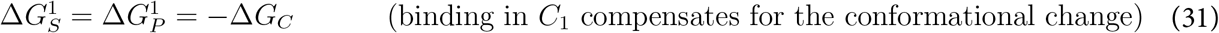

where equalities can be relaxed to differences of order RT, the dynamics is effectively described by

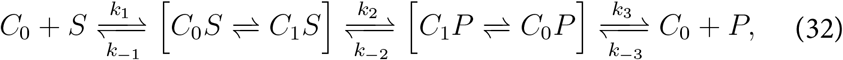

where the states within brackets are not distinguished, and define two effective states *CS* and *CP* (Fig. 3C). Formally, the kinetics is then equivalent to that describing single-state catalysis in Eq. (4).

In enzymes, the compensation between Δ*G*_*C*_ and 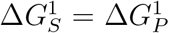required for Eq. (31) to hold can for instance take the form of an enthalpy-entropy compensation [52] between a high-entropy “open” state *C*_0_*S* where *S* is loosely bound to a flexible conformation *C*_0_ of the catalyst, and a high-enthalpy “closed” state where *S* is tightly bound to a rigid conformation *C*_1_ of the catalyst, in which case Δ*G*_*C*_ represents an entropic cost. Alternatively, or additionally, Δ*G*_*C*_ can represent a desolvation free energy from a solvated conformation *C*_0_ to a desolvated conformation *C*_1_ [53].

As we now show, it is precisely in the conditions described by Eqs (29)-(30)-(31) where the kinetics of two-state catalysis is formally equivalent to that of single-state catalysis that the presence of two underlying states makes an essential difference. While Eq. (4) applies in both cases, the way in which the kinetic rates depend on intrinsic parameters are not the same, and the trade-offs at play are radically different.

#### Two-state uniform binding

With single-state catalysts, we saw that uniform binding cannot confer catalysis by itself but can improve on a pre-existing catalytic mechanism by decreasing 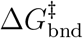at the expense of 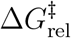, which is valuable when the substrate concentration is low (Fig. 2). With two state catalysts, uniform binding within each state cannot confer catalysis either, but, as we now show, it can improve on a pre-existing catalytic mechanism in the opposite case where release is limiting, by decreasing 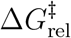at the expense of 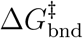, which is valuable when the substrate concentration is high.

This is achieved under the assumptions of Eqs. (29)-(30)-(31) that lead to an effectively unidimensional catalytic process with *N* = 2 states described by Eq. (32) and Fig. 3C. Under these assumptions the only free intrinsic parameter is Δ*G*_*C*_ *≥* 0. This parameter modifies, the intrinsic 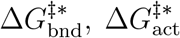 and 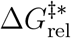of a pre-existing catalytic mechanism into (Fig. 4).

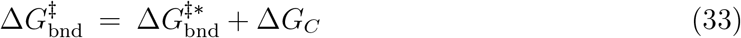

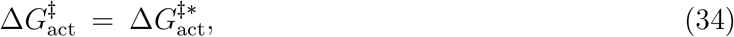

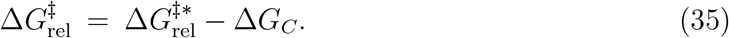

We thus obtain that Δ*G*_*C*_*≥* 0 plays exactly the same role as the uniform binding energy Δ*G*_*u*_ *≤* 0 for a one-state catalyst (Eq. (17) and Fig. 2), except that it has opposite sign and therefore opposite effects (Fig. 4): it lowers the release barrier at the expense of the binding/activation barrier 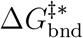. Provided 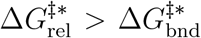, which occurs for sufficiently high substrate concentrations, a two-state mechanism is therefore advantageous, with an optimal value of Δ*G*_*C*_ given by 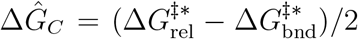. Furthermore, while uniform binding can only lower *k*_cat_ in the context of a single-state catalyst, it can increase it in the context of a two-state catalyst, since *k*_cat_ depends on 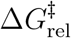 but not on 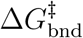. This is an example of a possibility that a conformational change offers beyond what rigid catalysts can possibly achieve. However, in this scenario as in Albery and Knowles’ original scenario [24], 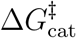 remains unchanged, and a pre-existing catalytic mechanism must be assumed for any catalysis to take place.

**FIG. 4:**
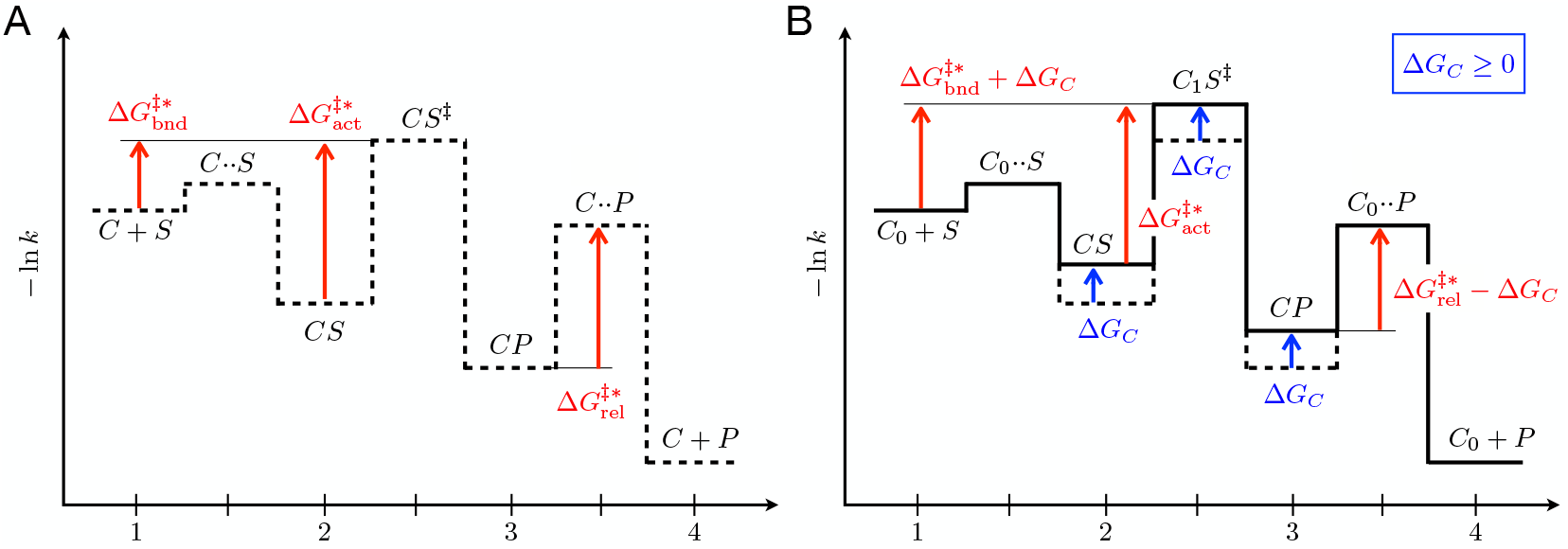
Two-state uniform binding. **A**. A pre-existing catalytic mechanism is assumed where 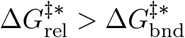, a situation opposite to Fig. 2A. **B**. Under the conditions of Fig. 3C with the further assumption that *C*_1_ binds uniformly to all reaction states, i.e., 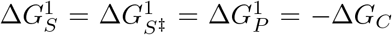 the only designable parameter is Δ*G*_*C*_ *>* 0, which can be chosen to have 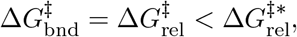, thus effectively reducing the cycling time *T*_*c*_. It does not affect, however, 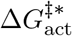which, as in the case represented here, may remain the limiting barrier.

#### Two-state differential binding

For a single-state catalyst, we saw that the constraint of differential binding sets a lower bound on the cycling time of the form 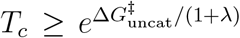, which excludes, in particular, perfect catalysis. As can be shown analytically and numerically (SM, Sec. 5 and Fig. S3D), the same bound applies to a two-state catalyst when each of its states is subject to the same constraint of differential binding, i.e., 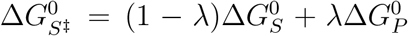 and 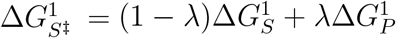. Under such constraints, the presence of two states cannot alleviate the fundamental limitations of single-state catalysts.

#### Two-state discriminative binding

In contrast, under constraints of discriminative binding where, in each state of the catalyst, arbitrary specificity to the transition state can be achieved at the expense of tight binding to the substrate and product, a two-state catalyst can overcome the limitations of single-state catalysis. Formally, the constraints of Eq. (22) are extended to two-state catalysts by imposing 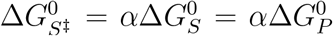 and 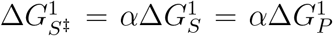. Catalytic “perfection” can even be reached (Fig. S3E). This is again achieved under the assumptions of Eqs. (29)-(30)-(31) that lead to an effectively unidimensional catalytic process with *N* = 2 states described by Eq. (32) and Fig. 3C. These assumptions leave only one designable parameter, namely Δ*G*_*C*_ *≥* 0. As illustrated in Fig. 5 for the case 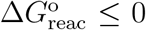, choosing this parameter to satisfy 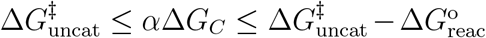, i.e., if the reaction is irreversible 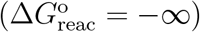,

**FIG. 5:**
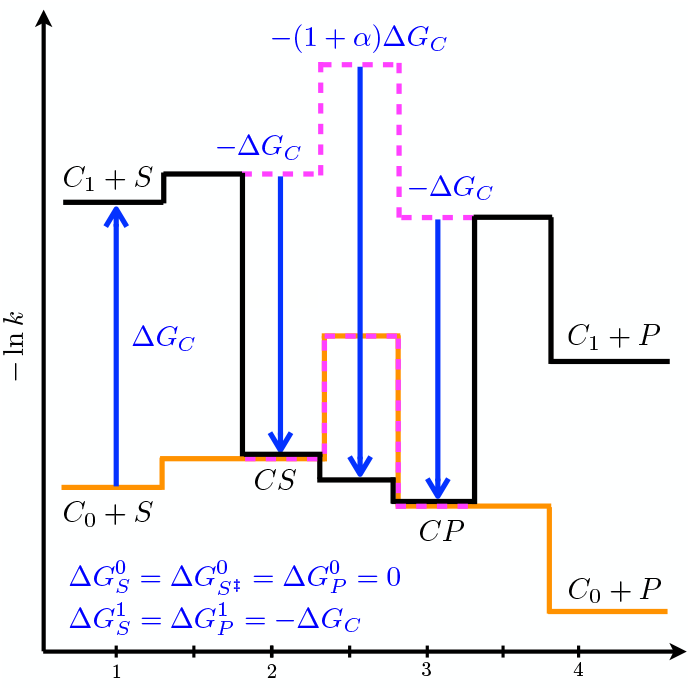
Perfect catalysis with two-state discriminative binding. We consider as in Fig. 3C a design verifying the conditions of Eqs. (29)-(30)-(31) so that the interconversions *C*_0_*S* ⇌ *C*_1_*S* and *C*_0_*P* ⇌*C*_1_*P* are instantaneous and define two effective states *CS* and *CP*. Under constraints of discriminative binding, the difference 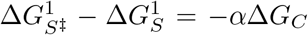can take arbitrary low values provided Δ*G*_*C*_ is large enough. A value of Δ*G*_*C*_ can thus be chosen so that *C*_1_*S → C*_1_*P* is barrier less. In cases where 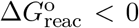, as illustrated here, this leaves, as the only kinetic barrier, the barrier associated with the diffusion of the substrate towards the catalyst, *C*_0_ + *S → C*_0_*S*. Perfect catalysis is then achieved that is only diffusion-limited.

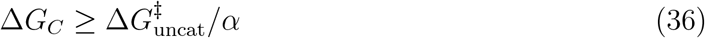

makes negative all barriers along the path *C*_0_ + *S → C*_0_*S → C*_1_*S → C*_1_*P → C*_0_*P → C*_0_ + *P*, except for the inevitable extrinsic barrier associated with diffusion at the first step *C*_0_ + *S → C*_0_*S*. Further, no state outside of this path is a kinetic trap: *C*_1_ +*S* relaxes to *C*_0_ +*S* without kinetic barrier and similarly for 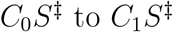and *C*_1_ + *P* to *C*_0_ + *P* .

This design can be understood as decoupling the activation and release steps, which are in trade-off in the other scenarios: activation is made to occur in one state of the catalyst – the active state *C*_1_ with a large binding free energy 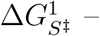while product release is made to occur in a different state – the inactive state *C*_0_ with negligible binding free energy 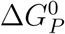. The switch between the two states is itself made barrier-less by introducing a large energy difference Δ*G*_*C*_ between *C*_0_ and *C*_1_ that compensates for 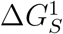and 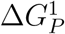, thus making the transitions *C*_0_*S → C*_1_*S* and *C*_1_*P → C*_0_*P* barrier-less. By this mechanism, Sabatier principle is abolished and perfect catalysis reached despite constraints of discriminative binding within each state of the catalyst. We previously illustrated this principle in a simple physics model [39] where we assumed 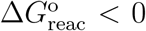, but it applies more generally to spontaneous reactions with arbitrary values of 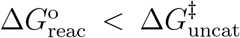, including cases where 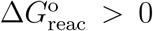, in which case Eq. (36) must be replaced by 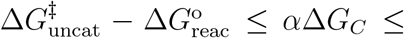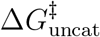, and perfect catalysis can be limited by the thermodynamical barrier 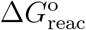when this barrier exceeds the diffusion barrier *−* ln(*k*_*d*_[*S*]) (SM, Sec. 3).

## CONCLUSION

Following and extending previous works by Albery and Knowles [24, 30], we analyzed the principles underlying enzymatic activities by treating catalysis as a modulation of kinetic barriers under constraints on the capacity to discriminate transition states from substrates and products. In the absence of such discrimination, unimolecular reactions cannot be catalyzed [14], but Albery and Knowles proposed that adding a non-discriminative interaction to a pre-existing catalytic mechanism was the predominant mechanism by which enzymes outperform chemical catalysts made up of small molecules [24, 30]. They further noted that such “uniform binding” was readily evolutionarily accessible through interactions with non-reactive “handles” that are part of many biological reactants. They contrasted this form of uniform binding with “differential binding”, where the affinity to the transition state is constrained to be intermediate between the affinities to the substrate and to the product, as commonly observed in chemistry [25, 49]. Here, we revisited this constraint of differential binding to demonstrate that it sets an upper bound on catalytic efficiency which excludes “perfect” catalysis, where rate acceleration is only limited by thermodynamics and diffusion. We pointed out that this limitation stems from the same trade-off between activation and release that is widely observed in heterogeneous catalysis where it underscores Sabatier principle of optimal catalysis [54].

To explain how enzymes can escape this trade-off and possibly reach perfection, we extended the model in two ways. First, we proposed that enzymes are better understood as subject to another form of constraints, which we called discriminative binding, where arbitrary specificity to the transition state is achievable but at the expense of increasingly large affinities to the substrate and product. This constraint formalizes the notion that high specificity to the transition state requires precise and rigid positioning of the substrate, which is possible only through strong interactions with non-reactive parts of the reactant that are common to the substrate and product. Second, we extended the analysis to catalysts that can be in several states, with different affinities to reactants in their different conformational states. This formalizes the observation that many enzymes undergo conformational changes and have catalytic activities that depend on their conformation, a property generally associated with allostery [21]. Our main conclusion is that two-state catalysts can overcome the limitations of single-state catalysts when subject to constraints of discriminative binding, but we also showed that two-state catalysts can exploit uniform binding to achieve the opposite effect of single-state catalysts, namely facilitating product release at the expense of a weaker enzyme-substrate association.

Our results demonstrate how conformational changes can play an essential role in catalysis, given constraints from chemical similarity between reaction states alone. This is to be contrasted with explanations for conformational changes in enzymes that refer to other types of constraints, e.g., constraints from substrate specificity, as in Koshland’s induced fit model [19, 20], constraints from geometry, as in models where a conformational change allows the enzyme to enclose a substrate without compromising its binding and release [15, 55], or constraints from regulation, as in many justifications of allostery [21]. This is also to be contrasted with proposals where conformational changes accelerate the chemical step through rate-promoting vibrations [56]. Our model is based on a definition of catalytic efficiency that takes into account the complete catalytic cycle, the role of a conformational change being to make the optimization of the chemical step *CS → CP* compatible with optimization of the subsequent step of product release *CP → C* + *P*. Our model is therefore consistent with rigid active sites being optimal for the chemical step [7, 8].

The two-state architecture that we find conducive to perfect catalysis is peculiar, with a weakly interacting state coexisting with a strongly interacting state of higher free energy. This architecture echoes the description of many enzymes as switching between an entropy rich inactive state and a rigid active state [57], a feature that has directly been observed in single molecule experiments [58]. The model also sets constraints on the free energy cost of the conformational change, which must be commensurate with the activation energy of the spontaneous reaction. To be precise, conformational changes are not strictly necessary to achieve the effects that our model describes: what matters is primarily a free energy difference between two states of the substrate-enzyme complex, which, for instance, may also be achieved through a distortion of the substrate.

The presence of two states with different affinities for a same ligand is a prominent feature of allostery [38]. Allostery, however, usually involves an effector that is distinct from the substrate and that binds at a site remote from the active site [21]. A parallel is made by viewing the substrate as made of two pieces, a reactive part and a non-reactive part, which bind to distinct – although generally not remote – sites of the catalyst, a binding site and an active site. In this “split-site” model [33], the non-reactive substrate handle acts as an allosteric effector. This is a very different role than in Albery and Knowles’ mechanism of uniform binding where the substrate handle acts as an entropic trap. For triosephosphate isomerase, the enzyme on which Albery and Knowles built their analysis, but also for several other enzymes, Richard and collaborators experimentally cut substrates in two pieces and showed that the dissociated non-reactive handles indeed act as allosteric effectors [10, 59]. Recognizing that transitions between conformational changes can play an essential role in catalysis independently of any regulation is consistent with the notion of latent allostery, the presence of cooperative effects preceding the evolution of allosteric regulation [60]. It also suggests an evolutionary scenario in which the coupling between chemical reactions and mechanical motion at work in molecular motors first appeared as a consequence of selective pressures on catalytic efficiency.

The view of substrate handles as enabling multi-state catalysis is closely related to Jencks’s proposal that these handles enable the expression in the transition state of an “intrinsic binding energy” which is only partially realized in the substrate-catalyst complex [9]. Our model may in fact be seen as a formalization of this proposal. This formalization provides at least three clarifications. Firstly, our model identifies the constraints under which this mechanism is necessary, namely the chemical similarity between reaction states. Secondly, our model links this mechanism to conformational changes and allostery, and thus provides a rationale for the prevalence of these features in enzymes. As noted previously, other mechanisms can possibly achieve the same effects. Jencks, in fact, downplayed the contribution of conformational changes [61] while emphasizing the role of substrate destabilization [26], but our model indicates that a comparable destabilization of the product is necessary. In our model, not all the forms of destabilization are as conducive to catalysis. In particular, a destabilization stemming from a physical distortion of the bound substrate that is released in the transition state as well as in the product state, which may be considered the “most obvious mechanism of substrate destabilization” [26], cannot achieve perfect catalysis unless another mechanism is present that destabilizes the bound product (SM Sec. 6).

We also noted that optimal two-state catalysis can be kinetically indistinguishable from single-state catalysis, and described by the same Michaelis-Menten kinetics (see Eq. (32)), which can explain why the contribution of conformational changes to enzymatic catalysis is often overlooked. However, the impact of mutations that reduce binding affinity to substrate handles on kinetic parameters depends on the underlying mechanism. Uniform binding predicts that decreasing this affinity increases *k*_cat_ (or, if the activation barrier 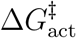dominates, that it leaves it nearly unchanged). Two-state catalysis under constraints of discriminative binding predicts, on the other hand, that *k*_cat_ decreases when the activation barrier dominates. This latter scenario is in agreement with many observations [9, 12]. Uniform binding and two-state catalysis are, however, non exclusive, and can even be complementary: uniform binding in the inactive state of a two-state enzyme can indeed provide the same benefits as uniform binding in a single-state enzyme under conditions of low substrate concentrations, by trading a slower release for a more efficient substrate capture. A role for conformational changes in catalysis is also not excluding other roles concomitantly played by the same conformational change, e.g., a role in enclosing the substrate and/or enabling regulation of the enzyme activity. Different scenarios have, however, important differences from an evolutionary perspective. Optimal uniform binding requires a fine-tuned affinity to the handle that depends on substrate concentration, on properties of the spontaneous reaction and on the catalytic mechanism, while optimal two-state binding requires primarily an affinity that compensates for the cost of the conformational change with a value that is only loosely constrained by the activation free energy of the spontaneous reaction. Such a mechanism opens the possibility for an enzyme to adapt to catalyze a new reaction while preserving the same two-state mechanism if the new and old substrates share the same handle. This possibility is consistent with the repeated attachment of the same handles to many substrates, e.g., phosphate handles to metabolites [62], as well as with the concomitant reutilization of the same folds, e.g., TIM barrels [63], in enzymes catalyzing different reactions.

The approach that we followed to rationalize enzyme mechanisms focuses on the constraints imposed by chemical similarity between reaction states. The importance of these constraints is well-recognized in heterogeneous catalysis, where they take the form of Sabatier principle [54]. This qualitative principle states that an optimal catalyst must strike a compromise between high affinities that lower the activation energy and low affinities that favor product release. Our analysis recovers this trade-off when the catalyst is single-state, whether the constraints take the form of differential binding or discriminative binding. Chemical constraints have been particularly studied for transition-metal catalysis, where they are found to follow scaling relationships, with a few “descriptors” linearly controlling the binding affinity of the catalytic surface to the different reaction states when comparing surfaces made of different metallic elements [42, 48]. In the context of the unimolecular reaction that we studied, this corresponds to the observation that transition-state and product affinities are both linearly related to substrate affinity, i.e., Δ*G*_*S‡*_ = *a*_*S‡*_ Δ*G*_*S*_ and Δ*G*_*P*_ = *a*_*P*_ Δ*G*_*S*_ with factors *a*_*S‡*_*≥* 0 and *a*_*P*_ *≥*0 that depend on the geometry of the surface but relate surfaces made of different metals. Formally, such scaling relations encompass uniform binding when *a*_*S‡*_ = *a*_*P*_ = 1, pure substrate stabilization when 1 *< a*_*S‡*_ = *a*_*P*_ (SM Sec. 6), differential binding when 1 *< a*_*S‡*_ *< a*_*P*_, and discriminative binding when 1 = *a*_*P*_ *< a*_*S‡*_. As we have shown, transitions between states allow for perfect catalysis in this later case, but also, more generally, whenever *a*_*P*_ *≤* 1 *< a*_*S‡*_ (SM Sec. 7). In the light of our model, the implementation of multiple states could overcome some of the limitations currently encountered in heterogeneous catalysis, but also in the design of new enzymes [64] where transitions between states is not currently envisaged.

## Acknowledgments

The author thanks Yann Sakref for comments, and ANR-21-CE45-0033, ANR-22-CE06-0037 for funding.

