## Supplementary material for "A role for conformational changes in enzyme catalysis"

Olivier Rivoire

Gulliver, CNRS, ESPCI, Université PSL, Paris, France.

#### 1. Cycling time $T_c$ as a function of elementary rates $k_{\pm i}$

Here we derive Eqs. (5)-(6)-(9) recursively by considering the mean first-passage time from $C+S$ to $C+P$ through a unidimensional Markov chain with $N$ intermediate states [44].

Starting with $N=1$, we consider a catalytic process with a single intermediate state, of the form $C+S \underset{k_{-1}}{\overset{k_{+1}}{\rightleftharpoons}} CS \underset{k_{-2}}{\overset{k_{+2}}{\rightleftharpoons}} C+P$. The time spent in the starting state $C+S$ before any transition is exponentially distributed with mean $1/k_1$ while the mean time spent in $CS$ is $1/(k_{-1}+k_2)$. Once in $CS$, the probability to transition to $C+P$ is $p=k_2/(k_{-1}+k_2)$. Given that this transition is made, the mean time from $C+S$ to $C+P$ is therefore $\tau=1/k_1+1/(k_{-1}+k_2)$. With probability $(1-p)$, however, a back transition is made to $C+S$, which takes additional time. If the system goes back to $C+S$ only once, which occurs with probability $(1-p)p$, the mean time from $C+S$ to $C+P$ is $2\tau$ and, more generally, if it goes back $n$ times, which occurs with probability $(1-p)^n p$, it is $(n+1)\tau$. In total, we therefore have $T_c=\sum_{n=0}^{\infty}p(1-p)^n(n+1)\tau=\tau/p$, which yields $T_c=1/k_1+1/k_2+k_{-1}/(k_1k_2)$. This result can be interpreted as the mean first-passage time for a Markov chain with no backward transition, of the form $C+S \xrightarrow{k_1'} CS \xrightarrow{k_2} C+P$, when considering an effective rate $k_1'$ defined by $1/k_1'=(1+k_{-1}/k_2)/k_1$, so that $T_c=1/k_1'+1/k_2$.

This interpretation is convenient to generalize the expression for $T_c$ to an arbitrary number $N$ of intermediate states. For the case of $N=2$ intermediate states that corresponds to Eq. (4), we can thus write $T_c=1/k_1'+1/k_2'+1/k_3$ with $1/k_1'=(1+k_{-1}/k_2')/k_1$ and $1/k_2'=(1+k_{-2}/k_3)/k_2$, which leads to

$$T_c=\frac{1}{k_1}+\frac{1}{k_2}+\frac{1}{k_3}+\frac{k_{-1}}{k_1k_2}+\frac{k_{-2}}{k_2k_3}+\frac{k_{-1}k_{-2}}{k_1k_2k_3}. \tag{S1}$$

This formula extends to $N$ intermediate states along a unidimensional chain of transitions,

$$T_c=\sum_{j=1}^{N+1}\sum_{j=i}^{N+1}\frac{1}{k_j}\prod_{r=i}^{j-1}\frac{k_{-r}}{k_r}=\sum_{1\leq i\leq j\leq N+1}\frac{k_{-i}k_{-(i+1)}\dots k_{-(j-2)}k_{-(j-1)}}{k_ik_{i+1}\dots k_{j-1}k_j}. \tag{S2}$$

This result can be obtained by several other approaches, including by writting $T_c=\sum_{i=1}^{N}1/k_i'$ where the times $1/k_i'$ are recursively defined by $1/k_{N+1}'=1/k_{N+1}$ and $1/k_i'=(1+k_{-i}/k_{i+1}')/k_i$ for $1\leq i\leq N$, with $1/k_i'$ representing the mean residence time in state $i$ [65].

Introducing $G_i$ and $G_i^\ddagger$ related to $k_{\pm i}$ by $k_i=e^{-(G_i^\ddagger-G_i)}$ and $k_{-i}=e^{-(G_i^\ddagger-G_{i+1})}$, Eq. (S2) can be rewritten as Eq. (9).

#### 2. Choice of intrinsic parameters

Single-state catalysis with $N=2$ intermediate states involves 6 rates, $k_{\pm1},k_{\pm2},k_{\pm3}$, of which only 5 are independent since the ratio $(k_1k_2k_3)/(k_{-1}k_{-2}k_{-3})$ is set by the equilibrium constant of the spontaneous reaction and the concentrations $[S]$ and $[P]$ of substrates and products. Out of these 5 parameters, 3 only are intrinsic, i.e., depend on properties of the catalyst, since $k_1 = k_D[S]$ and $k_2 = k_D[P]$ are set by the diffusion of substrates and products towards the catalyst.

One option is to retain as independent intrinsic parameters $k_{-1}, k_2, k_3$. We chose instead to take $\Delta G_S = -\ln k_{-1}$, $\Delta G_{S^\ddagger} = -\ln k_2 - \Delta G^\ddagger_{\text{uncat}}$ and $\Delta G_P = -\ln k_3$. Albery and Knowles [30] make a difference choice, and take as parameters $k_{-1}$, $K_{\text{int}}$ and $k_2^o$ where $K_{\text{int}} = k_2/k_{-2}$ and $k_2 = k_2^o K_{\text{int}}^\lambda$ with a given $\lambda$ (which they denote $\beta$).

Albery and Knowles take uniform binding to refer to the optimization of $T_c$ over $k_{-1}$ at fixed $K_{\text{int}}$ and $k_2^o$ [30]. We take it to refer to the optimization of $T_c$ over $\Delta G_u$ with $\Delta G_S = \Delta G_{S^\ddagger} = \Delta G_P = \Delta G_u$, but since $-\ln k_{-1} = \Delta G_u$ in this case, and since neither $k_2$ nor $k_{-2}$ depend on $\Delta G_u$, optimizing over $k_{-1}$ is equivalent to optimizing over $\Delta G_u$. The only difference that we introduce is a distinction between an uncatalyzed activation barrier with $-\ln k_2 = \Delta G^\ddagger_{\text{uncat}}$ and a catalyzed activation barrier with $-\ln k_2 < \Delta G^\ddagger_{\text{uncat}}$ that we associate with a pre-existing catalytic mechanism. This distinction is important for recognizing that uniform binding does not confer catalysis by itself.

Albery and Knowles take differential binding to refer to the optimization of $T_c$ over $K_{\text{int}}$ at fixed $k_{-1}$ and $k_2^o$ [30]. We take it to refer to the joint optimization of $T_c$ over $\Delta G_S$ and $\Delta G_P$ under the constraint that $\Delta G_{S^\ddagger} = (1-\lambda)\Delta G_S + \lambda\Delta G_P$. This is equivalent to a joint optimization over $K_{\text{int}}$ and $k_{-1}$ at fixed $k_2^o = e^{-\Delta G^\ddagger_{\text{uncat}}}$. Since $\ln K_{\text{int}} = -\Delta G^{\text{o}}_{\text{reac}} + \Delta G_S - \Delta G_P$, varying $K_{\text{int}}$ is indeed equivalent to varying the difference $\Delta G_S - \Delta G_P$. Varying jointly $\Delta G_S$ and $\Delta G_P$, i.e., $k_{-1}$ and $K_{\text{int}}$, is justified under an evolutionary scenario where uniform binding occurs on faster evolutionary time scales than differential binding.

#### 3. Extension to reversible reactions

The main text presents for simplicity the case $\Delta G^{\text{o}}_{\text{reac}} = -\infty$ so that the reverse transitions $P \to S$ and $CP \to CS$ are impossible. More generally, we may consider a reversible reaction with a finite $\Delta G^{\text{o}}_{\text{reac}}$ satisfying

$$\Delta G^{\text{o}}_{\text{reac}} < \Delta G^\ddagger_{\text{uncat}}, \tag{S3}$$

a condition that is necessary for the two states $S$ and $P$ to be well defined. When considering catalysis, another condition is for $CS^\ddagger$ to be a transition state, which takes the form

$$\Delta G^\ddagger_{\text{uncat}} + \Delta G_{S^\ddagger} \geq \max(\Delta G_S, \Delta G^{\text{o}}_{\text{reac}} + \Delta G_P). \tag{S4}$$

When considering reversible reactions, a fourth kinetic barrier is present that controls the cycling time $T_c$ in addition to the three kinetic barriers $\Delta G^\ddagger_{\text{act}}$, $\Delta G^\ddagger_{\text{rel}}$, $\Delta G^\ddagger_{\text{bnd}}$ defined in Eq. (14), namely

$$\Delta G^\ddagger_{\text{back}} = \Delta G^{\text{o}}_{\text{reac}} - \Delta G_S, \tag{S5}$$

which can be interpreted as arising from the possible recrossing of the transition state.

This additional barrier is also controlling $k_{\text{cat}}$ and the condition for catalysis $k_{\text{cat}} \geq k_0$ therefore requires $\Delta G^\ddagger_{\text{lim}} < \Delta G^\ddagger_{\text{uncat}}$ where

$$\Delta G^\ddagger_{\text{lim}} = \max(\Delta G^\ddagger_{\text{act}}, \Delta G^\ddagger_{\text{rel}}, \Delta G^\ddagger_{\text{back}}). \tag{S6}$$

The two conditions of Eq. (15) must then be extended to include a third condition,

$$\Delta G_S > \Delta G^{\text{o}}_{\text{reac}} - \Delta G^\ddagger_{\text{uncat}}. \tag{S7}$$

By itself, this third condition does not introduce any new trade-off: minimizing $\Delta G_{S^\ddagger}$ and maximizing $\Delta G_S$ and $\Delta G_P$ contribute to minimize each barrier.

In particular, the condition for $CS^\ddagger$ to be well defined, given by Eq. (S4), is not restrictive. If $\Delta G^{\text{o}}_{\text{reac}} < 0$, the optimal catalyst is only limited by diffusion, i.e., $\Delta G^\ddagger_{\text{lim}} = \Delta G^\ddagger_{\text{diff}}$, where

$$\Delta G^\ddagger_{\text{diff}} = -\ln(k_D[S]) \tag{S8}$$

represents the kinetic barrier for the diffusion of the substrate towards the catalyst. This is achieved by taking $\Delta G_P = 0$ and $\Delta G^{\text{o}}_{\text{reac}} \leq \Delta G_{S^\ddagger} + \Delta G^\ddagger_{\text{uncat}} \leq \Delta G_S \leq 0$. If $\Delta G^{\text{o}}_{\text{reac}} > 0$, the optimal catalyst is either limited by diffusion or by $\Delta G^{\text{o}}_{\text{reac}}$, which is achieved by taking $\Delta G_S = 0$ and $0 \leq \Delta G_{S^\ddagger} + \Delta G^\ddagger_{\text{uncat}} \leq \Delta G^{\text{o}}_{\text{reac}} + \Delta G_P$. In any case, we have a lower bound on the limiting barrier given by

$$\Delta G^\ddagger_{\text{lim}} \geq \max(\Delta G^\ddagger_{\text{diff}}, \Delta G^{\text{o}}_{\text{reac}}), \tag{S9}$$

where the limit value $\Delta G^\ddagger_{\text{lim}} = \max(\Delta G^\ddagger_{\text{diff}}, \Delta G^{\text{o}}_{\text{reac}})$ is in principle achievable and represents "perfect" catalysis. When the reaction is irreversible, perfect catalysis is equivalent to diffusion-limited catalysis but when $\Delta G^\ddagger_{\text{diff}} < \Delta G^{\text{o}}_{\text{reac}}$, perfect catalysis may rather correspond to "thermodynamics-limited catalysis". Because a finite $\Delta G^{\text{o}}_{\text{reac}}$ only adds a kinetic barrier, the bounds on $k_{\text{cat}}$ obtained when $\Delta G^{\text{o}}_{\text{reac}} = -\infty$ all apply to reversible reactions.

The case of single-state uniform binding with finite $\Delta G^{\text{o}}_{\text{reac}}$ is treated in the next section. For two-state uniform binding, the trade-off between $\Delta G^{\ddagger *}_{\text{rel}} - \Delta G_C$ and $\Delta G^{\ddagger *}_{\text{bnd}} + \Delta G_C$ presented in the main text when $\Delta G^{\text{o}}_{\text{reac}} = -\infty$ becomes at finite $\Delta G^{\text{o}}_{\text{reac}}$ a trade-off between $\max(\Delta G^{\ddagger *}_{\text{rel}}, \Delta G^{\ddagger *}_{\text{back}}) - \Delta G_C$ and $\Delta G^{\ddagger *}_{\text{bnd}} + \Delta G_C$, with an optimal value of $\Delta G_C$ obtained when the two terms in trade-off are the same, i.e.,

$$\Delta \hat{G}_C = \frac{\max(\Delta G^{\ddagger *}_{\text{rel}}, \Delta G^{\ddagger *}_{\text{back}}) - \Delta G^{\ddagger *}_{\text{bnd}}}{2}. \tag{S10}$$

This optimum is of value for sufficiently high concentrations of substrate, namely when $\Delta G^{\ddagger *}_{\text{bnd}} < \max(\Delta G^{\ddagger *}_{\text{rel}}, \Delta G^{\ddagger *}_{\text{back}})$ (Fig. 4).

For two-state discriminative binding with a finite $\Delta G^{\text{o}}_{\text{reac}}$, Eq. (36) must be generalized to

$$\min(0, -\Delta G^{\text{o}}_{\text{reac}}) \leq \alpha \Delta G_C - \Delta G^\ddagger_{\text{uncat}} \leq \max(0, -\Delta G^{\text{o}}_{\text{reac}}) \tag{S11}$$

to ensure that no kinetic barrier is introduced by lowering too much $CS^\ddagger$. When $\Delta G^{\text{o}}_{\text{reac}} > 0$, catalysis is perfect but not barrier-less since $C_0{\cdot\cdot}P$ has a free energy higher than $C_0{\cdot\cdot}S$.

#### 4. Single-state uniform binding

The most restrictive constraint is to assume single-state uniform binding controlled by a single degree of freedom $\Delta G_u \leq 0$ that forces all binding free energies to be the same,

$$\Delta G_S = \Delta G_{S^\ddagger} = \Delta G_P = \Delta G_u. \tag{S12}$$

By itself, uniform binding cannot confer catalysis [14], as it is apparent from the expression for $\Delta G^\ddagger_{\text{act}}$ in Eq. (14) which implies $\Delta G^\ddagger_{\text{act}} = \Delta G^\ddagger_{\text{uncat}}$ and therefore $k_{\text{cat}} \leq k_0$. It can, however, be beneficial in the context of a pre-existing mechanism [24]. To study this case, we consider that $\Delta G_X = \Delta G^*_X + \Delta G_u$ for each state $X = S, S^\ddagger, P$, where the $\Delta G^*_X \leq 0$ encode a pre-existing catalytic mechanism.

In this case, the four kinetic barriers controlling $T_c$ (considering here the general case of reversible reactions with $\Delta G^\ddagger_{\text{back}}$ defined in Eq. (S5)) are

$$\begin{aligned} \Delta G^\ddagger_{\text{bnd}} &= \Delta G^{\ddagger *}_{\text{bnd}} + \Delta G_u \\ \Delta G^\ddagger_{\text{act}} &= \Delta G^{\ddagger *}_{\text{act}} \\ \Delta G^\ddagger_{\text{rel}} &= \Delta G^{\ddagger *}_{\text{rel}} - \Delta G_u \\ \Delta G^\ddagger_{\text{back}} &= \Delta G^{\ddagger *}_{\text{back}} - \Delta G_u, \end{aligned} \tag{S13}$$

(S14)

where $^*$ refers to the pre-existing mechanism. As $k_{\rm cat}$ depends on the last three barriers only, it cannot be increased by $\Delta G_u \leq 0$. The first barrier, however, can be decreased. A decrease of $k_{\rm cat}$ can therefore be compensated by a decrease of $K_M$ to yield an overall lower $T_c$. The trade-off is between $\max(\Delta G^{\ddagger *}_{\rm rel}, \Delta G^{\ddagger *}_{\rm back}) - \Delta G_u$ and $\Delta G^{\ddagger *}_{\rm bnd} + \Delta G_u$ (Fig. 2). The optimal $\Delta G_u$ is obtained when the two terms in trade-off are the same, which corresponds to

$$\Delta \hat{G}_u = \frac{\max(\Delta G^{\ddagger *}_{\rm rel}, \Delta G^{\ddagger *}_{\rm back}) - \Delta G^{\ddagger *}_{\rm bnd}}{2}. \tag{S15}$$

This is relevant when $\Delta \hat{G}_u < 0$, for instance when the substrate concentration is sufficiently low to have $\Delta G^{\ddagger *}_{\rm bnd} > \Delta G^{\ddagger *}_{\rm rel}$ (Fig. 2).

This is the mechanism proposed by Albery and Knowles [24]. Our derivation differs from theirs by approximating $T_c$ with the largest (limiting) barrier. Not making this approximation, we can minimize $T_c$ over a rate $k_u = e^{\Delta G_u}$ that such that $k_{-1} = k^*_{-1} k_u$ and $k_3 = k^*_3 k_u$. The expression for $T_c$ is then

$$y = \frac{1}{k_1} + \frac{1}{k_2} + \frac{1}{k^*_3 k_u} + \frac{k^*_{-1} k_u}{k_1 k_2} + \frac{k_{-2}}{k_2 k^*_3 k_u} + \frac{k^*_{-1} k_{-2}}{k_1 k_2 k^*_3}. \tag{S16}$$

Differentiation with respect to $k_u$ leads to an optimum given by

$$k_u^2 = \frac{k_1 k_2}{k^*_3 k^*_{-1}} \left(1 + \frac{k_{-2}}{k_2}\right) \tag{S17}$$

when the right-hand side is $< 1$. It can be rewritten as Eq. (12) in [24] (with $\theta = 0$),

$$\frac{k_{-1}}{k_1 k_2} = \frac{1}{k_3} \left(1 + \frac{k_{-2}}{k_2}\right) \tag{S18}$$

which is equivalent with our notations to

$$e^{-\Delta G^{\ddagger}_{\rm bnd}} = e^{-\Delta G^{\ddagger}_{\rm rel}} + e^{-\Delta G^{\ddagger}_{\rm back}}. \tag{S19}$$

Retaining only the largest kinetic barrier, this corresponds to the criterion $\Delta G^{\ddagger}_{\rm bnd} = \max(\Delta G^{\ddagger}_{\rm rel}, \Delta G^{\ddagger}_{\rm back})$ by which we obtained the optimal value of $\Delta G_u$ in Eq. (S15).

### 5. Limits to two-state differential binding

A necessary (but not sufficient) condition for a two-state catalyst to outperform a single-state catalyst is that the two-dimensional network of transitions described in Eq. (27) contains a unidimensional path from $C_0 + S$ to $C_0 + P$ for which $T_c$ is lower. As explained in Fig. S1, out of the 8 possible paths that one can trace, only one can possibly provide a lower $T_c$, the path described by Eq. (28). We therefore show here that when considering this unidimensional Markov chain and assuming the constraint of differential binding in both states, $\Delta G^0_{S^\ddagger} = (1-\lambda)\Delta G^0_S + \lambda \Delta G^0_P$ and $\Delta G^1_{S^\ddagger} = (1-\lambda)\Delta G^1_S + \lambda \Delta G^1_P$, the same bound $\ln T_c \geq \Delta G^{\ddagger}_{\rm uncat}/(1+\lambda)$ that limits single-catalyst applies.

To this end, we consider in turn two cases, $\Delta G_C + \Delta G^1_S \leq 0$ or $\Delta G_C + \Delta G^1_S \geq 0$. Starting with $\Delta G_C + \Delta G^1_S \leq 0$, the two barriers $G_{C_1 S^\ddagger} - G_{C_0 S}$ and $G_{C_0 \cdot\cdot P} - G_{C_1 P}$, corresponding to the terms $G^\ddagger_3 - G_2$ and $G^\ddagger_5 - G_4$ in Eq. (9), are each a lower bound on $\ln T_c$. Since

$$\begin{aligned} G_{C_1 S^\ddagger} - G_{C_0 S} &= \Delta G^\ddagger + (\Delta G_C + \Delta G^1_S) - \Delta G^0_S \geq \Delta G^\ddagger_{\rm uncat} + (\Delta G_C + \Delta G^1_S), \\ G_{C_0 \cdot\cdot P} - G_{C_1 P} &= -(\Delta G_C + \Delta G^1_P), \end{aligned} \tag{S20}$$

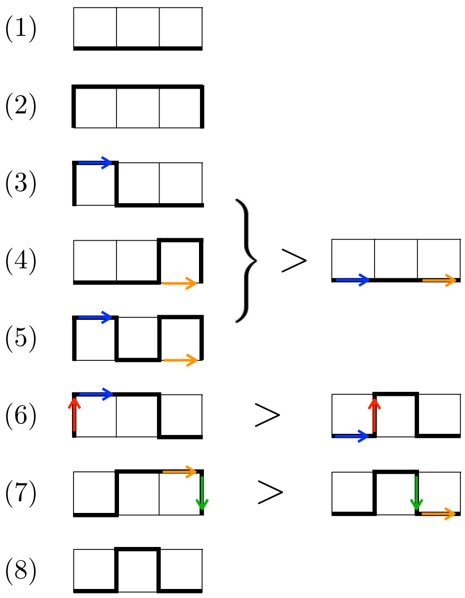


FIG. S1: Paths connecting $C_0+S$ to $C_0+P$ in the two-dimensional network of transitions described by Eq. (27). Only path (8) can possibly improve on the linear path of a single-state catalyst when considering constraints of differential binding. Path (1) is equivalent to a linear path and path (2) only adds new transitions to a linear path. Path (3) includes the same barrier $\Delta G^\ddagger_{\rm diff}$ as the linear path (blue arrow) and the detour that it provides cannot therefore improve on the linear path. Similarly for the detour provided by path (4) since the barrier $-\Delta G^0_P$ remains included in the expression for $T_c$ as $G^\ddagger_5 - G_3$ in Eq. (9) ($G_3 = G_{C_0P}$ and $G^\ddagger_5 = G_{C_0\cdot\cdot P}$). *A fortiori*, path (5) which has the two detours of paths (3) and (4) cannot improve on the linear path. Paths (6) and (7) are at best as efficient as path (8). This is because any benefit for going through $C_1S \to C_1P$ must come from the fact that $\Delta G^1_{S^\ddagger} - \Delta G^1_S < \Delta G^0_{S^\ddagger} - \Delta G^0_S$ which, under constraints of differential (or discriminative) binding, is only possible if $\Delta G^1_S < \Delta G^0_S$ and $\Delta G^1_P < \Delta G^0_P$. For path (6), the barriers associated with the blue arrows are same, namely $\Delta G^\ddagger_{\rm diff}$, while the barrier associated with the red arrows is $\Delta G_C + \Delta G^\ddagger_C$ for path (6) compared to $\Delta G_C + \Delta G^\ddagger_C + \Delta G^1_S - \Delta G^0_S < \Delta G_C$ for the linear path. For path (7), the barriers associated with the orange arrows are $-\Delta G^1_P$ to be compared to $-\Delta G^0_P < -\Delta G^1_P$ for the linear path and the barriers associated with the green arrows are $\Delta G^\ddagger_C$ in both cases. In conclusion, the most favorable path differing from the linear path is path (8), which justifies that we focus on the unidimensional Markov chain described by Eq. (27).

we have two lower bounds that depend on $\Delta G_C + \Delta G^1_S$ and $\Delta G_C + \Delta G^1_P$ in the same way as $\Delta G^\ddagger_{\rm act}$ and $\Delta G^\ddagger_{\rm rel}$ depend on $\Delta G_S$ and $\Delta G_P$ in Eq. (19), with the same constraints $\Delta G^\ddagger_S = (1-\lambda)\Delta G_S + \lambda \Delta G_P$ and $\Delta G_S \leq 0$ (from the assumption $\Delta G_C + \Delta G^1_S \leq 0$). The same lower bound $\Delta G^\ddagger/(1+\lambda)$ therefore applies in this case.

Next, we consider the case $\Delta G_C + \Delta G^1_S \geq 0$. The two barriers $G_{C_1S^\ddagger} - G_{C_0+S}$ and $G_{C_0\cdot\cdot P} - G_{C_1P}$, corresponding to the terms $G^\ddagger_3 - G_1$ and $G^\ddagger_5 - G_4$ in Eq. (9), are each a lower bound on $\ln T_c$. They satisfy

$$\begin{aligned} G_{C_1S^\ddagger} - G_{C_0+S} &= \Delta G^\ddagger_{\rm uncat} + \Delta G_C + (1-\lambda)\Delta G^1_S + \lambda \Delta G^1_P, \\ G_{C_0\cdot\cdot P} - G_{C_1S} &= -(\Delta G_C + \Delta G^1_P). \end{aligned} \tag{S21}$$

Given $\Delta G^1_S$, the largest of these two free energy is minimized by taking

$$\Delta G_C + \Delta \hat{G}^1_P = -\frac{\Delta G^\ddagger + (1-\lambda)(\Delta G_C + \Delta G^1_S)}{1+\lambda}. \tag{S22}$$

The barriers are then the same and decreasing with $\Delta G_S^1$. Given the assumption that $\Delta G_C + \Delta G_S^1 \geq 0$, the optimum is for $\Delta G_C + \Delta G_S^1 = 0$ which yields again the same lower bound $\ln T_c \geq \Delta G^\ddagger_{\rm uncat}/(1+\lambda)$. We therefore conclude that the presence of two states cannot overcome the limitations that the constraints of differential binding set on the cycling time.

#### 6. Single-state substrate destabilization

We examine here a scenario of catalysis by pure substrate destabilization, where the substrate is destabilized but not the product, which we formalize by assuming that

$$\begin{aligned} \Delta G_S &= \Delta G_d + \Delta G_u, \\ \Delta G_{S^\ddagger} &= \Delta G_P = \Delta G_u, \end{aligned} \quad \text{(S23)}$$

where $\Delta G_d \geq 0$ represents a destabilization that applies only to the substrate and where $\Delta G_u \leq 0$ represents a stabilization that applies to all reaction states, with $\Delta G_d + \Delta G_u \leq 0$ to guarantee that the substrate-catalyst complex $CS$ is well-defined (Fig. S2). In other words, $\Delta G_u$ represents an "intrinsic binding energy" that is not fully expressed in the enzyme-substrate complex [9]. In this case, the four intrinsic barriers controlling the cycling time are

$$\begin{aligned} \Delta G^\ddagger_{\rm bnd} &= \Delta G^\ddagger_{\rm diff} + \Delta G^\ddagger_{\rm uncat} + \Delta G_u \\ \Delta G^\ddagger_{\rm act} &= \Delta G^\ddagger_{\rm uncat} - \Delta G_d \\ \Delta G^\ddagger_{\rm rel} &= -\Delta G_u \\ \Delta G^\ddagger_{\rm back} &= \Delta G^{\rm o}_{\rm reac} - \Delta G_d - \Delta G_u. \end{aligned} \quad \text{(S24)}$$

$k_{\rm cat}$, which depends only on the last three barriers, is maximized by maximizing $\Delta G_d$ and minimizing $\Delta G_u$, which implies that the optimum verifies $\Delta G_d = -\Delta G_u$. The trade-off between $\Delta G^\ddagger_{\rm act} = \Delta G^\ddagger_{\rm uncat} - \Delta G_d$ and $\Delta G^\ddagger_{\rm rel} = \Delta G_d$ is then optimized by taking $\Delta \hat{G}_d = \Delta G^\ddagger_{\rm uncat}/2$, which leads to the bound

$$k_{\rm cat} \leq e^{-\Delta G^\ddagger_{\rm uncat}/2}. \quad \text{(S25)}$$

This bounds is reached when $\Delta G^\ddagger_{\rm diff} \to 0$ and $\Delta G^{\rm o}_{\rm reac} \to -\infty$, showing that substrate destabilization can provide catalysis, but not perfect catalysis. The fact that substrate destabilization as formalized by Eq. (S23) can lower activation barriers by at most a factor two is also verified in numerical simulations (Fig. S3B).

#### 7. Catalysis under constraints described by scaling relationships

In heterogeneous catalysis, scaling relationship are documented that have the form $G_X = a_X G_0 + b_X$ where $a_X \geq 0$ for each reaction state $X = S, S^\ddagger, P$ [42]. With $G_0 = 0$ describing no interaction, this corresponds to $\Delta G_X = a_X G_0$. Without loss of generality, we can take $G_S$ as a descriptor so that

$$\Delta G_{S^\ddagger} = a_{S^\ddagger} \Delta G_S \quad \text{(S26)}$$

$$\Delta G_P = a_P \Delta G_S \quad \text{(S27)}$$

where $a_{S^\ddagger} \geq 0$ and $a_P \geq 0$ are viewed as constraints and where $\Delta G_S \leq 0$ represents the only independent intrinsic parameter.

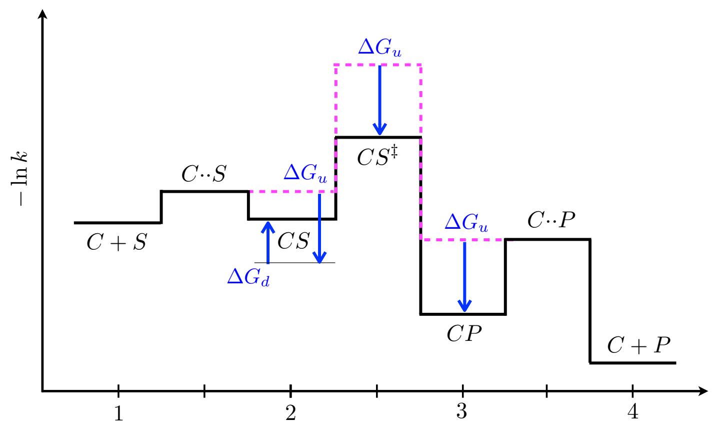


FIG. S2: Catalysis by substrate destabilization. All reaction states are subject to the same stabilizing free energy $\Delta G_u \leq 0$ and state $S$ is additionally subject to a destabilizing free energy $0 \leq \Delta G_d \leq -\Delta G_u$. This results in an activation barrier $\Delta G^\ddagger_{\rm act}$ lower than the activation barrier $\Delta G^\ddagger_{\rm uncat}$ for the spontaneous reaction (see Fig. 1) but at the expense of introducing a release barrier $\Delta G^\ddagger_{\rm rel} = -\Delta G_u$ which limits the efficiency of this mechanism.

In this context, we first consider single-state catalysts. The three kinetic barriers controlling $k_{\rm cat}$ have the form

$$\begin{aligned} \Delta G^\ddagger_{\rm act} &= \Delta G^\ddagger_{\rm uncat} + (a_{S^\ddagger} - 1)\Delta G_S \\ \Delta G^\ddagger_{\rm rel} &= -a_P \Delta G_S \\ \Delta G^\ddagger_{\rm back} &= \Delta G^{\rm o}_{\rm reac} - \Delta G_S. \end{aligned} \quad \text{(S28)}$$

Catalysis requires $\Delta G^\ddagger_{\rm act} < \Delta G^\ddagger_{\rm uncat}$ and therefore $1 < a_{S^\ddagger}$. We find again a trade-off between the activation barrier $\Delta G^\ddagger_{\rm act}$ and the other two, $\Delta G^\ddagger_{\rm rel}$, and $\Delta G^\ddagger_{\rm back}$. To obtain a simple bound, we can take the limit of irreversible reactions, $\Delta G^{\rm o}_{\rm reac} \to -\infty$, in which case the optimal trade-off is when $\Delta G^\ddagger_{\rm act} = \Delta G^\ddagger_{\rm rel}$, i.e., $\Delta \hat{G}_S = -\Delta G^\ddagger_{\rm uncat}/(a_S + a_P - 1)$. This implies

$$k_{\rm cat} \leq e^{\Delta G^\ddagger_{\rm uncat}/(1+\mu)} \qquad \text{with} \qquad \mu = \frac{a_{S^\ddagger} - 1}{a_P - 1}. \quad \text{(S29)}$$

The particular case $a_{S^\ddagger} < a_P$ is a form of differential binding with $\lambda = (a_{S^\ddagger} - 1)/(a_P - 1)$. Since $\lambda > \mu$, we verify that Eq. (S29) indeed implies Eq. (21).

When $a_P = 1$ in addition to $a_{S^\ddagger} > 1$, this is a particular instance of discriminative binding with $\alpha = a_{S^\ddagger} - 1$. We can then achieve perfect catalysis through a two-state allosteric catalyst. Other values of $a_P \neq 1$ may also allow it, the constraint on $a_P$ being that the transition $C_1P \to C_0P$ is barrier-less, along with the transition $C_0S \to C_1S$, i.e. $\Delta G_C + \Delta G^1_S \leq 0$ and $\Delta G_C + \Delta G^1_P = \Delta G_C + a_P \Delta G^1_S \geq 0$, which translates into the condition $a_P \leq 1$. Even when this condition is not satisfied, however, allostery may permit to overcome the limitations of Eq. (S29).

### 8. Numerical results

The bounds on the cycling time that we derive analytically can be verified through numerical simulations by estimating $T_c$ for systems whose kinetic rates are sampled at random. Fig. S3 shows results where the free energies are chosen uniformly at random within an interval, retaining only values for which the conditions given by Eqs. (S3) and (S4) are satisfied. In all cases, $\Delta G^\ddagger_{\rm diff} = -\ln(k_D[S]) \sim U([0, 10])$,

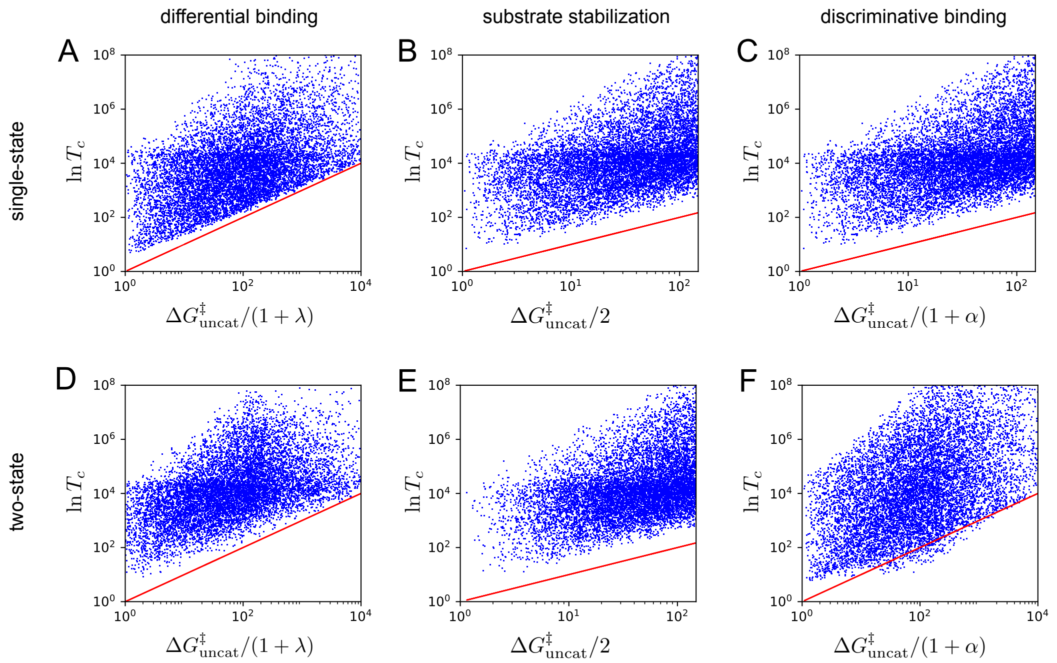


FIG. S3: Numerical results showing the cycling time $T_c$ of $10^4$ systems whose kinetic rates are drawn at random (see SM Sec. 8). **A.** Single-state differential binding, verifying $\ln T_c \geq \Delta G^{\ddagger}_{\rm uncat}/(1+\lambda)$ (blue points all above the red line).**B.** Single-state substrate destabilization, verifying $\ln T_c \geq \Delta G^{\ddagger}_{\rm uncat}/2$. **C.** Single-state discriminative binding, verifying $\ln T_c \geq \Delta G^{\ddagger}_{\rm uncat}/(1+\alpha)$. **D.** Two-state differential binding, verifying $\ln T_c \geq \Delta G^{\ddagger}_{\rm uncat}/(1+\lambda)$. **E.** Two-state substrate destabilization, verifying $\ln T_c \geq \Delta G^{\ddagger}_{\rm uncat}/2$. **F.** Two-state discriminative binding with an inactive state $C_0$, verifying that $\ln T_c$ can be lower than $\Delta G^{\ddagger}_{\rm uncat}/(1+\alpha)$ (some blue points are below the red line).

$\Delta G^{\ddagger}_{\rm uncat} \sim U([0,10])$, $\Delta G^{\rm o}_{\rm reac} \sim U([-10,10])$ and $\Delta G_S \sim U([-10,0])$ where $U([a,b])$ denotes a uniform distribution in the interval $[a,b]$. When considering differential binding (Fig. S3A,D), $\lambda \sim U([0,1])$, $\Delta G_P \sim U([0,1])$ and $\Delta G_{S^{\ddagger}} = (1-\lambda)\Delta G_S + \lambda G_P$. When considering substrate destabilization (Fig. S3B,E), $\Delta G_d \sim U([0,10])$. When considering discriminative binding (Fig. S3B,E), $\alpha \sim U([0,1])$, $\Delta G_P = \Delta G_S$ and $\Delta G_{S^{\ddagger}} = (1+\alpha)\Delta G_S$. When considering two-state catalysts, $\Delta G^{\ddagger}_C = 0$ and we either draw $\Delta G^0_S$ and $\Delta G^1_S$ each independently in $U([-10,0])$ (Fig. S3D) or take $\Delta G^0_S = 0$ (inactive $C_0$) and $\Delta G^1_S \sim U([0,1])$ (Fig. S3E,F). In all cases but with two-state discriminative binding (Fig. S3F), we verify the bounds $\ln T_c \geq \Delta G^{\ddagger}_{\rm uncat}/(1+\lambda)$ in the case of differential binding, $\ln T_c \geq \Delta G^{\ddagger}_{\rm uncat}/2$ in the case of substrate destabilization, and $\ln T_c \geq \Delta G^{\ddagger}_{\rm uncat}/(1+\alpha)$ in the case of discriminative binding. $T_c$ is computed through a matrix inversion using standard methods for the computation of mean first-passage times [39].
